# TIMEOR: a web-based tool to uncover temporal regulatory mechanisms from multi-omics data

**DOI:** 10.1101/2020.09.14.296418

**Authors:** Ashley Mae Conard, Nathaniel Goodman, Yanhui Hu, Norbert Perrimon, Ritambhara Singh, Charles Lawrence, Erica Larschan

## Abstract

Uncovering how transcription factors (TFs) regulate their targets at the DNA, RNA and protein levels over time is critical to define gene regulatory networks (GRNs) in normal and diseased states. RNA-seq has become a standard method to measure gene regulation using an established set of analysis steps. However, none of the currently available pipeline methods for interpreting ordered genomic data (in time or space) use time series models to assign cause and effect relationships within GRNs, are adaptive to diverse experimental designs, or enable user interpretation through a web-based platform. Furthermore, methods which integrate ordered RNA-seq data with transcription factor binding data are urgently needed. Here, we present TIMEOR (Trajectory Inference and Mechanism Exploration with Omics data in R), the first web-based and adaptive time series multi-omics pipeline method which infers the relationship between gene regulatory events across time. TIMEOR addresses the critical need for methods to predict causal regulatory mechanism networks between TFs from time series multi-omics data. We used TIMEOR to identify a new link between insulin stimulation and the circadian rhythm cycle. TIMEOR is available at https://github.com/ashleymaeconard/TIMEOR.git.

## Introduction

Cellular responses are a complex cascade of interacting factors, including many genes that influence each other at the DNA, RNA and protein levels, either directly or indirectly. These cascades become apparent when altering a specific regulatory mechanism through a biological experiment, with the end goal of constructing a directed *gene regulatory network* (GRNs), which provides a temporal model to define such cellular responses (Chai et al., 2014). Most interactions within GRNs are poorly understood, with many of the genes involved remaining unidentified (Sahraeian et al., 2017). Using multi-omics techniques, including RNA-seq (Wang et al., 2009) and protein-DNA interaction (e.g. CUT&RUN (Skene and Henikoff, 2017) and ChIP-seq (Johnson et al., 2007)) data, biologists measure changes in gene expression and transcription factor binding over time so as to accurately reconstruct GRNs (Brent, 2016). Specifically, a system is perturbed over time in ordered ‘case’ experiments and compared with the normal (or ‘control’) experiment(s) to determine the differentially expressed (DE) genes, that is, the set of genes showing statistically significant quantitative changes in expression levels between two experiments. These time series data are essential to accurately identify altered regulatory mechanisms and reconstruct GRNs.

To identify key regulators of a system of interest, raw multi-omics data must be preprocessed, followed by a DE analysis and finally GRN reconstruction. No end-to-end pipeline exists that given raw omics data produces a GRN. Current pipeline methods that incorporate only a subset of these steps suffer from two additional limitations, 1) they do not accurately process and determine DE of temporal data, nor 2) predict the order of action of the key regulators of transcription, i.e. *transcription factors* (TFs) (Sahraeian et al., 2017; Wani and Raza, 2019). While there are several pipelines to analyze each omic data type (RNA-seq or ChIP-seq) in isolation, joint analysis of multiple omic data types is required to uncover a GRN that models changes at different molecular levels and over time (Barbosa et al., 2018; Marbach et al., 2012). Finally, stand-alone GRN inference methods exist but do not fully leverage available data. In what follows we discuss current methodology in light of these limitations.

### Current pipeline methods are unable to effectively uncover altered regulatory mechanisms

To accurately process and determine differential expression for temporal data, it is key to understand that regulatory mechanisms often involve a ‘cause-and-effect’ relationship of two types. First, *intervention cause* in a system where we ask: Is there an effect upon change of input?, and second, *predictive* (i.e. Granger) *cause* for GRNs where we ask: Can we infer a lead and lag relationship between genes (Granger et al., 1969)? To address intervention cause, we need to perform a sequence of established steps to pre-process the RNA-seq data, by performing quality control, alignment, read count calculation, filtering, normalization and correction. Several pipeline methods codify these steps to enable users to subsequently perform DE analysis and summarize uncovered regulatory mechanisms (Afgan et al., 2018; Torre et al., 2018; Ge et al., 2018; Cornwell et al., 2018; de Jong et al., 2015; Jensen et al., 2017; Kartashov and Barski, 2015; Spurney et al., 2020).

However, there are many possible methods to choose from at each step in this process (**STAR Methods**), not all experimental designs are the same, and downstream results heavily depend on how the RNA-seq data are processed. Most current pipeline methods do not allow users to compare the many possible methods, providing only one tool for each step (BioJupies (Torre et al., 2018); RSEQREP (Jensen et al., 2017)) even though it is recommended to consider methods for analyses such as DE (Spies et al., 2019). Furthermore, most only perform part of the analysis to infer altered regulatory mechanisms (VIPER (Cornwell et al., 2018); BioWardrobe (Kartashov and Barski, 2015)). Also, the majority of platforms do not use multiple data-types (iDEP (Ge et al., 2018); T-REx (de Jong et al., 2015)), even though we know that RNA-seq does not provide direct evidence of gene interaction. Other pipeline methods used to process such RNA-seq data have to be pieced together such as Galaxy (Afgan et al., 2018) and do not use a time series DE model for time series data such as TuxNet (Spurney et al., 2020) thus disregarding time dependencies. **Figure S1** provides an overview of the available methods.

In fact, most pipeline methods used for RNA-seq data analysis use categorical DE methods that do not consider time dependency to determine which genes are interesting within time series data sets. Importantly, time series DE methods algorithms are still in their infancy and are underutilized (Spies et al., 2019). The few DE methods that do consider temporal dynamics are not yet part of pipeline methods to infer GRNs and are therefore incomplete (Fischer et al., 2018; Jensen et al., 2017; Michna et al., 2016; Nueda et al., 2014; Spurney et al., 2020; Wani and Raza, 2019). For time series DE methods that do exist, some can only analyze specific experimental designs or require normalized and corrected data on input (Sahraeian et al., 2017; Spies and Ciaudo, 2015; Spies et al., 2019).

After DE analysis, these pipeline methods aim to output regulatory mechanisms which include interacting genes. There are one or more of three main analysis outputs: 1) semi-informative gene ontology (GO) plots, showing what processes are affected and those genes involved; 2) pathways (i.e. directed gene graphs), showing directionality on a predefined network at a short time-scale with no crosstalk connections between pathways; and 3) networks (i.e. undirected gene graphs), thus losing directionality between genes and conflating gene and protein identity, yet gaining crosstalk between pathways and longer time scales. However, these three outputs do not include a GRN: a directed network highlighting how the key regulatory genes influence each other over time. It is important that pipeline methods enable users to compare multiple methods for RNA-seq processing steps to accommodate diverse experimental designs (Spies et al., 2019). Overall, no current pipeline method uses time series algorithms to measure gene dynamics over time and integrates transcription factor binding data to reconstruct GRNs as directed networks from ordered RNA-seq data.

### Current stand-alone GRN inference methods do not fully leverage available data

To predict the order of action of transcriptional regulators, there are several stand-alone methods which infer GRNs given processed data rather than raw sequence reads. These methods use time series and processed multi-omics data to establish which genes influence the expression of others downstream, providing information about the system dynamics (Mochida et al., 2018). Specifically, they address two main questions: 1) What genes are altered and exhibit dynamic expression changes?, 2) How are these genes within the GRN regulating each other? However, most of these stand-alone GRN inference methods only consider RNA-seq data from a single time point (i.e. at steady-state) (Langfelder and Horvath, 2008; Villaverde et al., 2014; Wang et al., 2013; Zhang et al., 2016; Huynh-Thu et al., 2010; Bar-Joseph et al., 2003; Faith et al., 2007; Haury et al., 2012; Liu et al., 2016; Margolin et al., 2006; Kang et al., 2018). While fewer methods consider consecutive time points (Huynh-Thu and Geurts, 2018; Bansal et al., 2006; Radovic et al., 2017; Roy et al., 2013; Zoppoli et al., 2010). Importantly, several consecutive time point GRN inference methods are limited in focus, constructing GRNs only when a gene of interest is present such as TSNI (Bansal et al., 2006) and may inaccurately interpret the network because they do not consider the experimental design. Thus, there is an urgent need for a pipeline method to leverage time series, known experimental design, gene interactions, and multi-omics data to provide evidence of interaction between genes, focusing on the master regulators of transcription, TFs, to reconstruct interpretable and high-fidelity GRNs in a user-guided fashion.

In summary, there is no integrated end-to-end analysis pipeline to accurately process and infer GRNs from temporal and multi-omics data. These deficiencies demand an adaptive, streamlined pipeline method that: a) supplies tailored methods for diverse experimental designs; b) performs multi-omics and experimentally determined protein-protein interaction data integration; c) uses time series models to generate informative predictions about altered regulatory mechanisms and GRNs; and d) provides an accessible and interactive interface to interpret multi-omics results.

### TIMEOR: new web-based tool to define regulated gene networks from multi-omics data

To address the critical gap to analyze and integrate time series RNA-seq and TF binding data, we present TIMEOR: Trajectory Inference and Mechanism Exploration using Omics data in R (**Figure 1**). This is the first automated interactive web (Chang et al., 2020; Sievert et al., 2020) (**Figure S2, S3**) and command line time series, multi-omics pipeline method for DE and comprehensive downstream analysis. TIMEOR reconstructs interpretable and user-guided GRNs. Specifically, TIMEOR’s web-interface leverages users’ knowledge of the biological process by providing options to users during each analysis phase. TIMEOR retrieves raw sequence reads (RNA-seq .fastq files) and performs all analysis from quality control and DE to enrichment (**Figure 1A, B**). Next, TIMEOR integrates motif and ChIP-seq data to define TF binding patterns, and reconstructs a TF GRN (**Figure 1C**). The web-interface enhances reproducibility and ease for users to dedicate more time to interpreting results and planning follow-up experiments (**Figure S2, S3**). Overall, TIMEOR is an adaptive method to predict GRNs from time series RNA-seq which integrates motif enrichment, ChIP-seq data, and temporal gene interactions.

**Figure 1:**
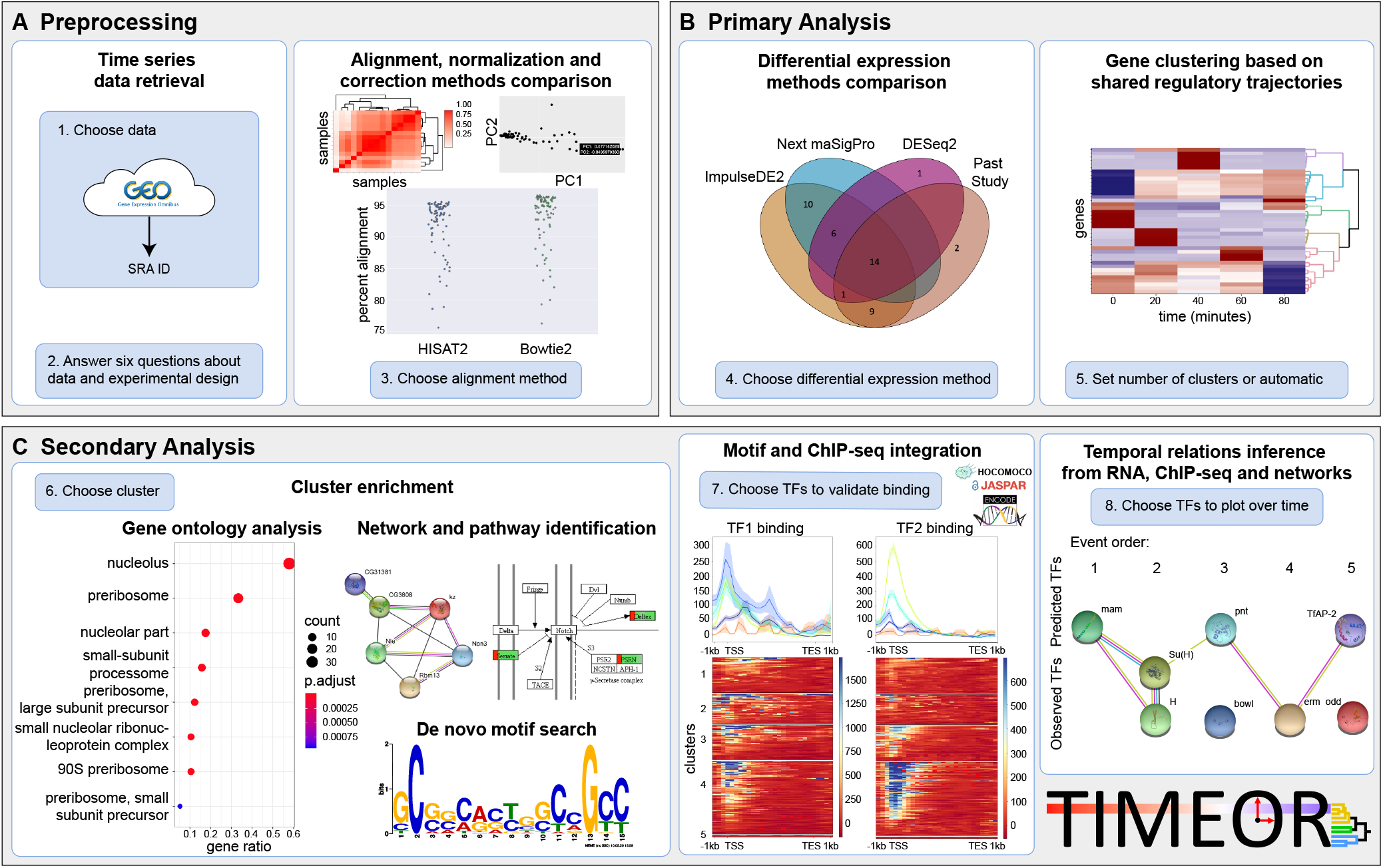
TIMEOR enables users to interrogate and reconstruct gene regulatory networks. Blue boxes denote main user-guided steps. **A.** Users specify a time series RNA-seq data set in the Pre-process Stage in which the data are automatically retrieved, normalized, corrected, filtered, and aligned to the selected reference genome using multiple alignment methods for comparison. **B.** The resulting count matrix after alignment is passed to Primary Analysis where multiple differential expression (DE) methods are run to produce a time series clustermap of DE over time. The user may also input the count matrix directly, skipping step A. **C.** DE results are passed to Secondary Analysis for gene ontology, pathway, network, motif, ChIP-seq analysis, and gene regulatory network construction.

We validated TIMEOR on both simulated and real data, demonstrating that TIMEOR can predict known and novel gene relationships within TF GRNs. Using real time series RNA-seq data collected after insulin stimulation (Zirin et al., 2019), TIMEOR discovered a novel ordered cascade of TF-TF interactions, providing a new link between insulin stimulation and a validated circadian clock GRN (Fathallah-Shaykh, 2010; Kadener et al., 2007; Maury, 2019; Zhou et al., 2016). TIMEOR revealed that insulin regulates several key circadian clock TFs, providing a new molecular mechanism linking a high sugar diet to sleep disruption (Qian and Scheer, 2016; Sharp et al., 2017; Stenvers et al., 2019). Overall, TIMEOR facilitates future analyses of integrated omics data to uncover novel biological and disease mechanisms from temporal gene expression data sets.

## Results

### The TIMEOR application package: accessible and streamlined tool to uncover novel regulatory mechanisms

The TIMEOR web-app (www.timeor.org) hosted at DRSC/TRiP (https://fgr.hms.harvard.edu/tools) and Harvard Medical School Research Computing, allows users to automatically and interactively analyze their time series data from *Drosophila melanogaster, Mus musculus, and Homo sapiens* (**Figure S2, S3**). The web-app and software package consists of multiple stages (**Figure 1**). First, the Pre-process Stage automatically chooses several methods to optimally preprocess the user data and generate results after the user answers six questions (**Figure 1A, S2A, S2B, S2C, S2D**). Second, in the Primary Analysis stage, DE results are compared between multiple continuous and categorical time series methods (Fischer et al., 2018; Nueda et al., 2014; Love et al., 2014). The user can also compare new results with a previous study to determine which DE method results to use for downstream analysis. TIMEOR then automatically clusters and plots (Sievert et al., 2020) the selected DE gene trajectories over time (**Figure 1B, S2D**). Third, the results are sent to the Secondary Analysis phase (**Figure 1C**) where three categories of analysis are performed in different tabs: *Enrichment*, identifies the genes and gene types that are over-represented within each cluster (**Figure S3A**); *Factor Binding*, predicts which TFs are post-transcriptionally influencing the expression of each gene cluster using motif and ChIP-seq data (**Figure S3B**); and *Temporal Relations*, identifies TFs GRN (**Figure S3D**). Each tab takes users through a series of exploratory results to determine the best predicted TF GRN (**STAR Methods**). In the following, we provide additional details on these three stages.

Overall, compared to current pipeline methods, TIMEOR has five unique features: 1) adaptive default analysis methods that can be customized to each experimental design; 2) multiple method comparisons for alignment and DE (for distant and close time point data); and 3) statistical, graphical and interactive results for data exploration. 4) Within each cluster of similarly regulated genes, TIMEOR performs automated gene enrichment, pathway, network, motif, and ChIP-seq analysis. 5) Lastly, TIMEOR merges experimentally determined gene networks (Franceschini et al., 2013), time series RNA-seq and motif and ChIP-seq information to reconstruct TF GRNs with directed causal interaction edges by labeling the causal interaction and regulation (activation or repression) between genes and gene products.

#### Pre-process Stage

The first stage of the TIMEOR package is the pre-process stage where users can load and retrieve published data sets of interest and perform preliminary analyses (**Figure 1A**). To do so, users can find their desired time series RNA-seq data on Gene Expression Omnibus (GEO) and upload GEO’s automatically generated file (Edgar et al., 2002). TIMEOR will then automatically generate a corresponding metadata file for subsequent analyses. After answering six basic questions regarding the experimental design, TIMEOR establishes the tailored adaptive default methods to use for RNA-seq data processing (**Figure S2A**). TIMEOR then automatically: 1) retrieves and stores the raw FASTQ files of interest in a unique directory created for each user; 2) checks RNA-seq data quality; 3) uses the adaptive defaults to compare and choose from several methods to align and calculate read counts per gene, resulting in a gene-by-replicate read count matrix (**Figure S2B**). TIMEOR provides several options to correlate and perform principal component analysis (PCA) between sample replicates, both automatically output as interactive plots. From these results, users can choose from several options to filter, normalize, and correct the resulting count matrix in preparation for the next stage (**Figure S2C, S2D**).

#### Primary Analysis

In Primary Analysis, users leverage the adaptive default methods to perform the most appropriate gene DE analysis on their data, or if the user answered “yes” to the question “Compare multiple methods (alignment/mapping, and differential expression)”, then multiple DE methods (continuous and categorical) are automatically run, and users can visualize the output in both a table and Venn diagram illustrating overlapping DE genes at a p-value threshold input by the user (Fischer et al., 2018; Nueda et al., 2014; Love et al., 2014). A previous study can be added to this Venn diagram to help the user determine which DE method results to use for downstream analysis. TIMEOR automatically clusters the chosen method’s DE gene trajectories, while also providing suggestions for manual cluster choice (**Figure 1B, S2E**). These cluster results are then passed to Secondary Analysis (**STAR Methods**).

#### Secondary Analysis

On the first tab of Secondary Analysis (**Figure 1C**) called *Enrichment*, users can toggle through each cluster to explore several types of enrichment within each cluster, including GO, pathway (Luo and Brouwer, 2013), network (Franceschini et al., 2013), and *de novo* motif discovery (Bailey and Elkan, 1994) (**Figure S3A**). On the second tab, *Factor Binding*, TIMEOR determines *observed TFs* (i.e. DE genes that are TFs) and *predicted TFs* (i.e. TFs enriched to bind to DE genes within a cluster) (**Figure S3B**). TIMEOR provides four predicted TF tables: 1) *Top Predicted Transcription Factors by Orthology* table, 2) *Top Predicted Transcription Factors by Motif Similarity* table, and their associated *Top Transcription Factors Table per Method* for both orthology and motif similarity (**Figure S3C**).

In order to identify predicted TFs and produce the first two tables described above, TIMEOR uses Rcistarget (Aibar et al., 2017) to scan up to 21 TF prediction methods for evidence of enriched TF binding to each cluster of genes by both *orthology* and *motif similarity* (**STAR Methods**). That is, each enriched motif is associated with candidate TFs based on orthologous sequences or based on similarities between annotated and unknown motifs (Aibar et al., 2017; Verfaillie et al., 2014). TIMEOR then summarizes the top predicted TFs for each TF prediction method and reports the *percent concordance* (i.e. the percent of methods that report the same most frequent TF) among these methods for the top four (default but changeable by user) predicted TFs. TIMEOR provides a consensus if more than 40% (default threshold but changeable by user) of the methods report the same top TFs. (**STAR Methods**) TIMEOR then scans ENCODE (Davis et al., 2018; Joly Beauparlant et al., 2020) for ChIP-seq data identifiers (IDs) for all predicted TFs. TIMEOR uses this ranking technique to output four predicted TF tables, that are distinguished by the type of evidence (orthology or motif similarity) and the method (consensus TF among all methods or a TF rankings within each method) (see **STAR Methods** and **Figure S3C**).

On the third tab, TIMEOR uses this observed and predicted TF information to determine the *Temporal Relations Table* between TFs, which defines a directed TF GRN (**Figure S3D**). By default, TIMEOR uses the *Top Predicted Transcription Factor by Orthology* table results to infer the GRN but the user is encouraged to consider all four predicted TF tables (**Figure S3C**). To characterize each temporal TF interaction within the GRN, we define a quadruple: (source TF, target TF, regulation type, interaction type) which we will explain next. *Interaction types* are: “predicted TF to observed TF, known interaction”; “predicted TF to observed TF, predicted interaction”; “observed TF to observed TF, known interaction”; or “observed TF to observed TF, predicted interaction”. *Regulation types* are “activation” or “repression” (**STAR Methods**). TIMEOR does this by using the temporal dynamics between clusters to assign known or predicted interaction edges between observed and predicted TFs observed within a window of time defined by users (question six **Figure S1A**). Additionally, for each *known* (i.e. experimentally determined) edge in STRINGdb (Franceschini et al., 2013), TIMEOR provides PubMed (https://www.ncbi.nlm.nih.gov/pubmed) identifiers to publications supporting that edge where applicable. Thus, using time series RNA-seq, motif and ChIP-seq analysis, and experimentally determined network information, TIMEOR generates an experimentally testable TF GRN (**Figure S3D**). TIMEOR provides a tutorial (**Figure S3E**) and outputs all results in the user’s personal analysis session folder to download for future use (**Figure S3F**).

### Evaluation of TIMEOR on Simulated Data

To determine TIMEOR’s robustness to recover TF GRNs, we simulated temporal expression patterns in *Homo sapiens* for genes known to interact with each other. Specifically, we used Polyester (Frazee et al., 2015) to simulate four RNA-seq expression cascading activation patterns over two biological replicates of six time points (**Figure S4E**) for 63 genes at approximately 20x sequencing coverage. We used the first time point as the control for the subsequent five time points. To simulate adequate background gene expression, we sampled and analyzed two biological replicates from a real time series RNA-seq experiment of fusobacterium nucleatum-stimulated human gingival fibroblasts control samples (GEO: GSE118691) taken at 2, 6, 12, 24, 48 hours (Kang et al., 2019a; Kang et al., 2019b). We simulated a constant RNA-seq expression of one fold across all six timepoints for all those genes that were active in at least one time point. To assess TIMEOR’s performance in inferring the ground truth GRN, we varied the percent concordance of the top TF identified by each method. For a concordance value of 35% and above, TIMEOR recovered simulated TF GRNs with perfect precision (**Figure 2A**). As expected, at low percent concordance between TF prediction methods TIMEOR predicts other TF interactions which were not simulated. Furthermore, TIMEOR achieved perfect recall for all percent concordance thresholds except at high percent concordance (65% and above) which lead to one ground truth TF to drop out. This simulation highlights TIMEOR’s ability to recover true temporal relations between TFs by integrating time series RNA-seq, motif data, and experimentally determined gene interaction information (**STAR Methods**).

**Figure 2:**
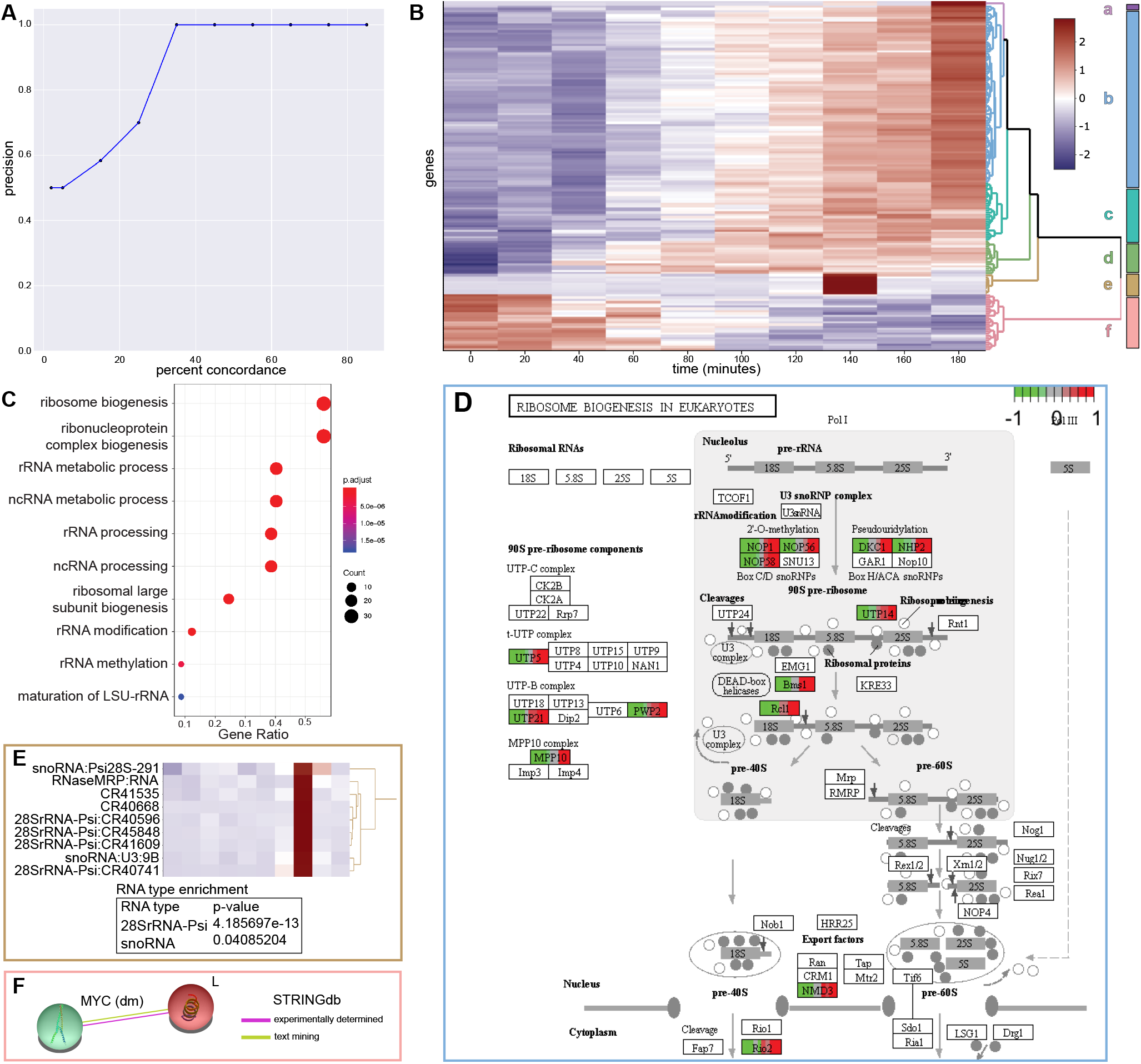
TIMEOR accurately recovers known and novel genomic relationships. **A.** Precision by percent concordance curve of TIMEOR’s ability to recover a simulated *Homo sapiens* TF GRN where the percent concordance between TF prediction methods ranges from 2% −85%. **B.** TIMEOR’s clustermap (Sievert et al., 2020) of significant genes’ trajectories using z-score to denote change in downregulation (blue) and upregulation (red). **C.** Without clustering, we recapitulate previous findings (Zirin et al., 2019) and illustrate these through GO analysis. **D.** TIMEOR’s pathway analysis for largest *cluster b*, which identifies the same ribosome biogenesis pathway as Zirin et al. 2019. **E.** TIMEOR output shows that in *cluster e* snoRNA pseudogenes and 28SrRNA pseudogenes are enriched (hypergeometric test). **F.** TIMEOR identified Lobe, which interacts directly with MYC (Wang and Huang, 2009). This interaction was not identified by prior analysis even though MYC was the focus of the previous study (Zirin et al., 2019).

### Evaluation of TIMEOR on Real Data

We next ran TIMEOR on insulin stimulation from Zirin et al., 2019 (Zirin et al., 2019), who performed an RNA-seq time series experiment where they incubated *Drosophila* SR2+ cells with insulin and performed RNA-seq on 10 consecutive time points every 20 minutes with three biological replicates. In the previous study with these data, Zirin et al. found that the MYC TF regulates tRNA synthetases which enhance growth of cells where MYC is overexpressed. Another class of MYC targets, ribosome biogenesis genes, are also affected in the time series RNA-seq data. Next, we describe how TIMEOR recapitulated previous findings and generated novel insights by analyzing each cluster separately (**Figure 2**), uncovered temporal dynamics among observed and predicted TFs by merging cluster information (**Figure 3**), and enabled us to formulate a new biological hypothesis regarding insulin stimulating the circadian rhythm cycle (**Figure 4**).

**Figure 3:**
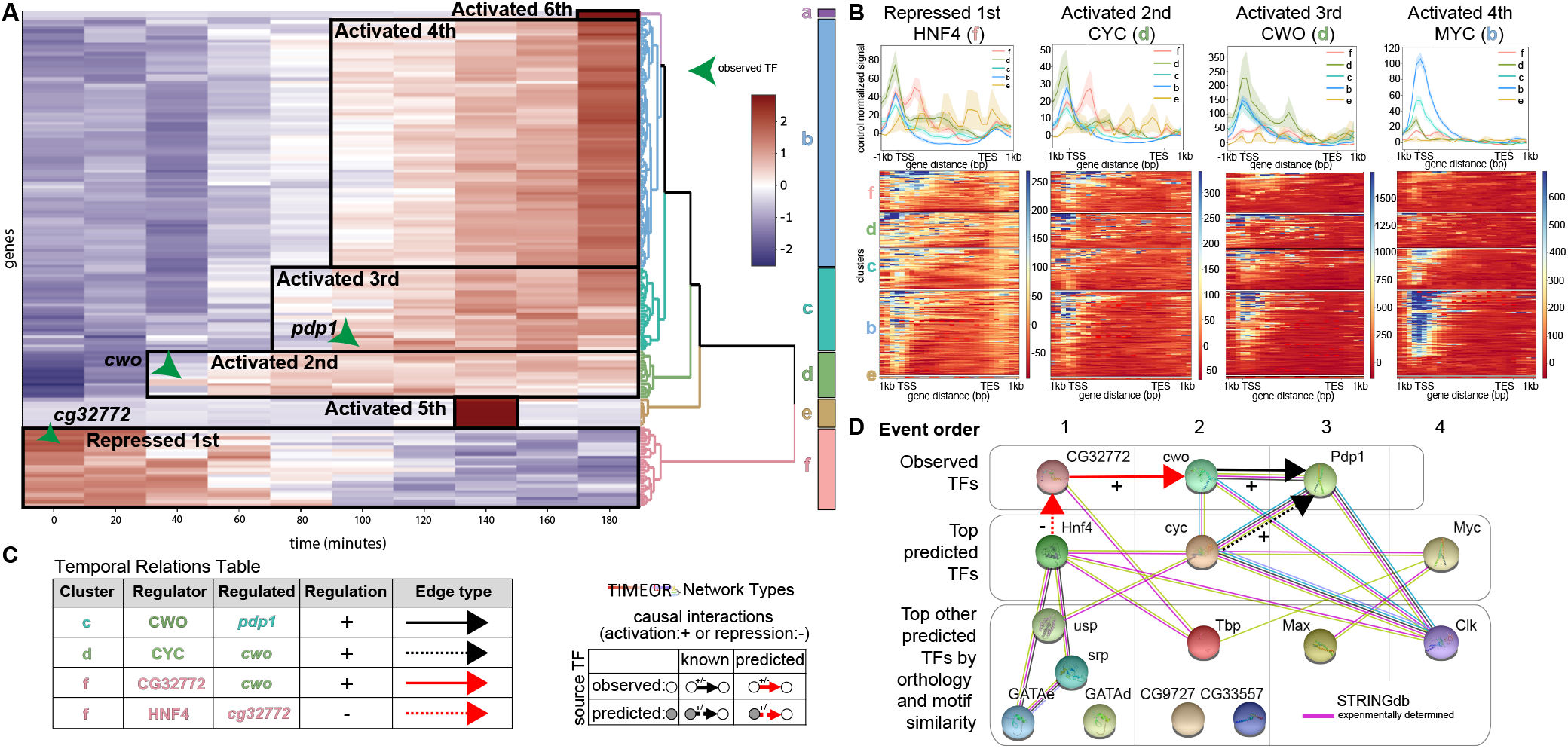
TIMEOR leverages temporal and multi-omics data to identify a novel gene regulatory network. **A.** TIMEOR’s clustermap (Sievert et al., 2020) highlighting TIMEOR’s temporal clustering with observed TFs present in the earliest changing clusters. **B.** Validation of predicted TFs using ChIP-seq data provided through TIMEOR to show binding affinity to genes across each cluster. **C.** Excerpt of TIMEOR’s Temporal Relations Table showing temporal relationship between TFs. **D.** Combining TIMEOR’s Temporal Relations Table, predicted TF information, and STRINGdb (Franceschini et al., 2013), TIMEOR outputs a gene regulatory network connecting insulin signaling and the circadian rhythm cycle.

**Figure 4.**
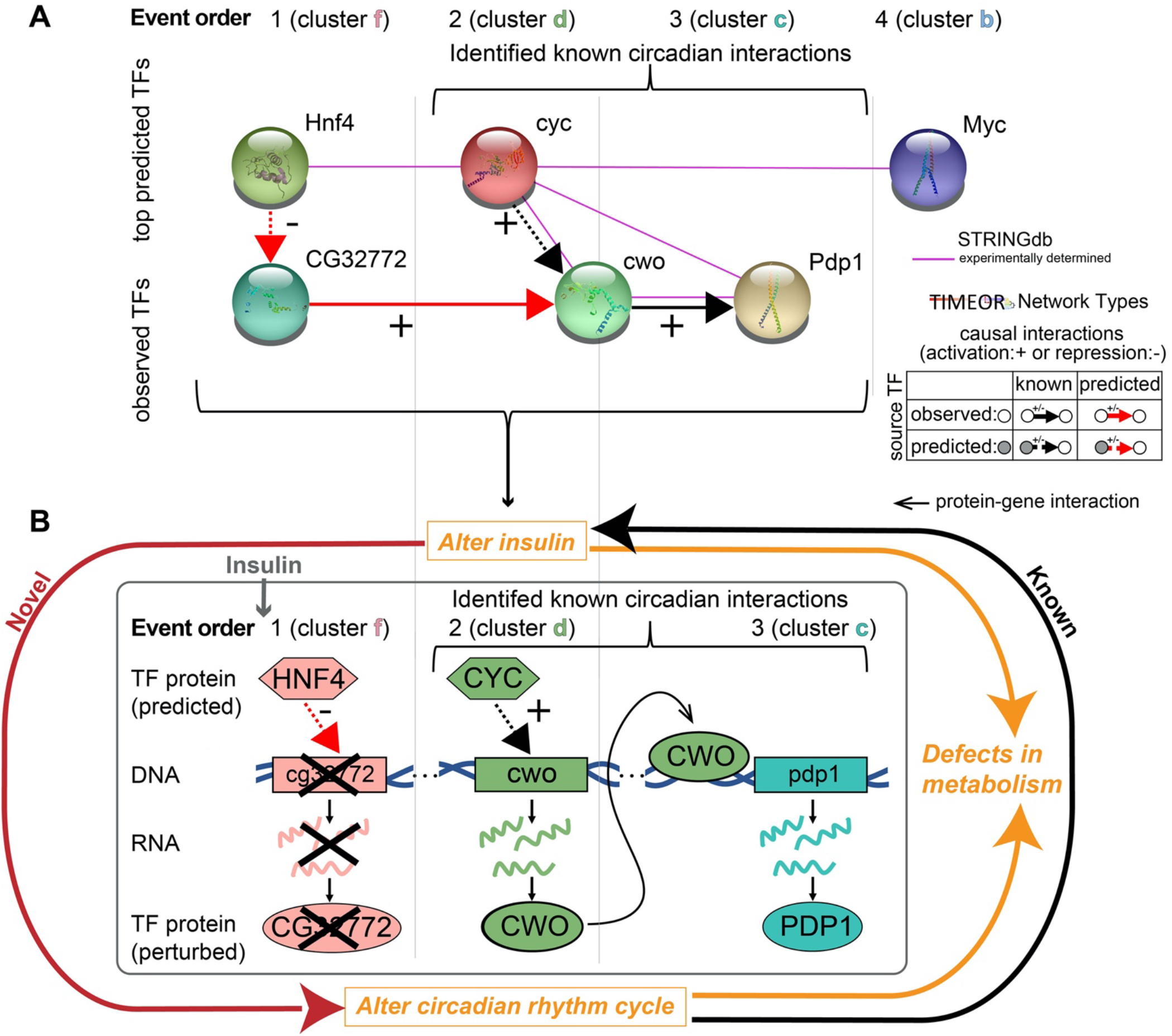
TIMEOR identifies that insulin acts as a cue for the circadian clock. **A.** TIMEOR identifies a GRN comprising both observed and the top predicted TFs binding each cluster (i.e. event order). The grey box denotes a known circadian rhythm causal interaction subnetwork (Fathallah-Shaykh, 2010; Zhou et al., 2016). **B.** Schematic showing that TIMEOR found two new players predicted to influence circadian rhythm cycle and are induced by insulin: HNF4, which is known to regulate glucose-dependent insulin secretion, and CG32772, which could regulate the *cwo* gene.

Using these time series data, TIMEOR recapitulates previous findings (Zirin et al., 2019) and discovers novel insights into insulin signaling. TIMEOR compared results from three different DE methods: DESeq2 (Love et al., 2014), Next maSigPro (Nueda et al., 2014), and ImpulseDE2 (Fischer et al., 2018). ImpulseDE2 showed the most significant overlap with the list of 1211 DE genes from Zirin et al. (p-value = 5.33342e-127 using the hypergeometric test) and the highest overlap with other methods (**Figure S4C**). Zirin et al. filtered and followed up on 33 of these 1211 genes to create a highly specific set. When TIMEOR overlapped these three methods with those 33 genes, ImpulseDE2 again showed the highest overlap with the previous study (p-value = 7.929393e-137 using the hypergeometric test) and other methods. We used GO analysis within TIMEOR to restate the findings from Zirin et al., 2019 (Zirin et al., 2019) which showed enrichment for genes regulating ribosome biogenesis before clustering the DE data (**Figure 2C**).

Next, TIMEOR clustered (Sievert et al., 2020) the DE data from ImpulseDE2 into six clusters using Euclidean distance between gene trajectories and Ward’s Method (Murtagh et al., 2014) to relate clusters of genes (**Figure 2B**). TIMEOR then generated the pathway, network, and GO analysis for each cluster. We found that the *cluster b*, the largest cluster, showed enrichment in the ribosome biogenesis (**Figure 2D**) pathway which was previously identified (**Figure 2C**). Moreover, TIMEOR’s clustering analysis (**Figure 2B**) showed that there were five additional clusters with different temporal patterns that were not identified by Zirin et al.

Importantly, TIMEOR determined that new information may be gained by considering the dynamics of gene expression patterns when performing time series experiments. TIMEOR found that the other gene trajectories represented by *clusters a, c, d, e*, and *f* each reveal new insights about insulin signaling (**Figure 2B**). *Clusters a* and *e* contained novel genes identified by TIMEOR that did not overlap with the previous study (Zirin et al., 2019): snoRNA and 28SrRNA pseudogenes. For these sets of noncoding genes, most pathway and GO analysis do not work, but TIMEOR is able to still demonstrate that these genes are enriched through a hypergeometric significance test (**Figure 2E**). These 28SrRNA-Psi may encode ribosomal RNA fragments that have been mislabeled as pseudogenes (Berk, 2016; Sharp, 1991, 2009). *Cluster a* only contained two genes, one snoRNA and one 28SrRNA pseudogene and interestingly these DEs followed a similar trajectory to *cluster e* genes that differed only late in the time series. *Cluster f* contains the group of genes that are repressed earliest in the regulatory cascade, including Lobe which was not identified as DE in the previous study even though it interacts directly with MYC (**Figure 2F**) (Wang and Huang, 2009).

The key task is to uncover how each gene cluster influences the expression of the other gene clusters through TFs by generating a GRN. To this end, TIMEOR integrates temporal dynamics between gene trajectory clusters with predicted TFs and STRINGdb’s experimentally determined network to temporally relate both observed (**Figure 3A arrows**) and predicted TFs (**Figure 3B, S3C**) within each cluster in a Temporal Relations Table (**Figure 3C**). TIMEOR identified the circadian rhythm TFs *pdp1* and *cwo* as observed in gene expression in response to insulin early in the regulatory cascade (**Figure 3A**). In the earliest regulated *cluster f* (repressed over time), TIMEOR identified the observed TF gene *cg32772*, followed by *cwo* in *cluster d* and *pdp1* in *cluster c*.

Next, TIMEOR leveraged motif analysis and ChIP-seq data to uncover and validate TF regulators for each cluster (**Figure 3B**). TIMEOR identified the available ENCODE data for all predicted TFs, as ChIP-seq data were not available after insulin stimulation. We chose the most comparable ChIP-seq data to input into TIMEOR to generate predicted TF binding profiles (**Figure 3B**). As predicted by the *Top Predicted Transcription Factors Table by Orthology*, Hnf4 showed the strongest binding affinity to the early repressed *cluster f* genes (gene body) and to the promoters of the next activated cluster (*cluster d* genes).

Next, we examined the two most popular TF prediction methods JASPAR (Sandelin et al., 2004) and HOMER (Heinz et al., 2010) and found that CYC ranks as the second most likely TF to bind *cluster d* (**Figure S4F**). As predicted, when examining TF binding, CYC bound most strongly to *cluster d* at the promoter. Moreover, CYC is known to bind to most of the observed TFs, and is known to bind to many of the other top predicted TFs (Franceschini et al. 2013; Fathallah-Shaykh, 2010; Zhou et al., 2016). CYC also showed strong binding within the gene body of genes in *cluster f*. CYC is likely the most enriched to bind *cluster d*.

CWO was observed in *cluster d* and bound strongly to the transcription start sites (TSS) of genes in that cluster (**Figure 3B**), as well as *cluster c* where there is a known interaction with *pdp1* (Fathallah-Shaykh, 2010; Zhou et al., 2016). In the next activated cluster, MYC showed strong binding to the later expressed genes in *cluster b* as predicted in the *Top Predicted Transcription Factors by Orthology* table. Therefore, average profiles in combination with TF prediction results (**Figure S4F**) provide support that the predicted TFs identified by TIMEOR are likely to be involved in regulating insulin signaling.

Importantly, *Cluster d* demonstrates how the user can leverage TIMEOR’s four predicted TF tables to identify temporal relationships between TFs (**Figure S4F**). Each TF prediction method is run individually on each cluster of genes, hence not integrating information across clusters. Thus, TIMEOR enables the user to view results across all clusters and generates new testable hypotheses through its four predicted TF tables and GRNs. The *Top Predicted Transcription Factors by Motif Similarity* table reported CG9272 as the most likely TF to bind *cluster d* and ChIP-seq results showed strong binding to the gene body of genes in *cluster d* (**Figure S4H**). In the *Top Predicted Transcription Factors by Orthology* table, TIMEOR reported TBP as the top predicted TF to bind (**Figure S4F**). However, TBP had an indistinguishable average binding profile across most clusters likely because it is a general TF (**Figure S4G**). Yet, there are *known* (i.e. experimentally determined) interactions between TBP and the observed TF CG32772 and TBP and the predicted TF Hnf4 in *cluster f* (Franceschini et al., 2013) (**Figure 3D**). Therefore, TIMEOR generated a testable hypothesis that TBP functions with CG32772 and Hnf4 to modulate gene regulation.

Integrating the observed and predicted TFs within TIMEOR’s Temporal Relations Table, TIMEOR augmented the experimentally determined STRINGdb network to predict a new GRN stimulated by insulin (**Figure 3D, 4**). That is, TIMEOR posited that the top predicted TF Hnf4 binding to *cluster f* repressed the observed gene *cg32772*. CG32772 was then predicted to activate *cwo* in the next activated cluster. Both of these interactions were predicted. CYC was predicted to bind genes in *cluster d* and activate the *cwo* gene in a *known* (i.e. experimentally determined) interaction. CWO is known to bind to *pdp1* in the next activated *cluster c*.

CYC, CWO and PDP1 are essential TFs regulating the circadian rhythm cycle and coordinate in the temporal order identified by TIMEOR (Fathallah-Shaykh, 2010; Zhou et al., 2016). These data support a model in which the TF CYC is binding to the *cwo* gene, which is then activated and influences the expression of *cluster c* (**Figure 4B**). Further, the CWO TF is found to bind to the gene encoding the observed TF *pdp1* in *cluster c*, and this information is thus highlighted by directed arrows in these networks (**Figure 4A and B**). Consistent with our predicted GRN, Zhou et al. (Zhou et al., 2016) and Fathallah-Shaykh et al. (Fathallah-Shaykh, 2010) illustrate that CYC functions earlier than CWO and *cwo* functions earlier than *pdp1* during the regulation of circadian rhythms.

In contrast to the circadian rhythm genes, little is known about observed TF CG32772 with the exception that it contains a zinc finger domain. However, given the temporal dynamics showed that CG32772 was repressed prior to the activation of the *cwo* gene, TIMEOR predicted that when repressed, CG32772 activates the expression of the observed *cwo* gene which is a novel and testable hypothesis. For example, the *CG32772* gene could be depleted to test its role in insulin signaling. Overall TIMEOR linked insulin signaling to a known circadian GRN and identified a new predicted TF that can be investigated in the future.

## Discussion

### Novel, accessible and reproducible pipeline method integrating time series RNA-seq data

Here, we presented Trajectory Inference and Mechanism Exploration with Omics data in R (TIMEOR). This pipeline method provides several advances including being the first accessible, adaptive and streamlined pipeline method to infer temporal dynamics between genes by intelligibly using multi-omics data, including time series RNA-seq and protein-DNA data (ChIP-seq or CUT&RUN) from multiple experimental designs. TIMEOR can compare multiple alignment and DE methods automatically and provide options for users to choose which results to use downstream. Our method provides multiple close and distant time point DE methods, and performs automatic gene trajectory clustering while also enabling users to manually cluster their data. The web-interface for TIMEOR provides users with the ability to interact, explore, and test variants of multiple types of analysis. Importantly, TIMEOR enables users to infer temporal relations between TFs and their gene targets through joint use of time series and multi-omics data and experimentally determined gene interaction networks.

TIMEOR enables researchers to run new analyses while also being able to reproduce results from past studies in a simple to use interface. Future work includes providing an RNA-seq read trimming method, and more alignment and DE methods for users (Bolger et al., 2014). Further, many comparative studies suggest taking a union of the DE results between methods for a more robust set of genes in downstream analysis, thus we would like to provide this functionality to the user (Spies et al., 2019). We plan to provide more interactive features such as integrating visNetwork (Thieurmel et al., 2016) to visualize networks, and integrate methods to examine temporal differential splicing and isoform expression. Furthermore, we would also like to add a set of methods specifically targeted for single-cell RNA-seq analysis. Lastly, there are several stand-alone GRN construction methods from time series RNA-seq data available, which we plan to integrate into TIMEOR. Our modular framework enables addition of novel and existing methods for pre-processing to GRN construction in future versions.

### Insulin acts as a cue for the circadian clock

We used TIMEOR to identify TF GRNs that are activated by insulin stimulation with two goals: 1) to uncover the dynamic changes in molecular function after insulin stimulation; 2) to identify the order of action of TFs in this process by integrating temporal RNA-seq data, ChIP-seq data, and experimentally determined gene interaction networks. Overall, TIMEOR recapitulated the findings from a previous study (Zirin et al., 2019) while also determining that insulin signaling activates the transcription of TFs that regulate the circadian rhythm cycle (**Figure 4A**). More research is needed to understand this relationship, although several studies suggest that the circadian clock coordinates the daily rhythm in human glucose metabolism (Boden et al., 1996; Stenvers et al., 2019).

Insulin resistance is a major contributor to the development of Type 2 Diabetes, caused by disrupting insulin, which alters metabolism (**Figure 4B**). The circadian rhythm cycle regulates several daily processes including metabolizing glucose to be used as energy and regulating insulin sensitivity (Qian and Scheer, 2016; Stenvers et al., 2019). Disrupting the circadian rhythm cycle (by staying up late or shift work) dysregulates metabolism, which can cause weight gain and type II diabetes (James et al., 2017). Therefore, it is known that disrupting the circadian clock can dysregulate insulin signaling. However, TIMEOR found a novel link, which shows that insulin directly regulates TFs that are known to alter the circadian clock such as *cyc*, *cwo*, and *pdp1*. TIMEOR found two new TFs that are predicted to influence the circadian rhythm cycle and are induced by insulin: 1) HNF4 is known to regulate glucose-dependent insulin secretion (Barry and Thummel, 2016); and 2) CG32772 could be regulating the *cwo* gene. Further validation experiments will be required to test these new hypotheses.

Overall, TIMEOR is a new adaptive method capitalizing on cause-and-effect modeling to analyze time series RNA-seq data and ChIP-seq. TIMEOR’s automated pipeline structure facilitates in depth analysis of input data while supporting reproducibility and providing the best tools for a given experimental design. Importantly, TIMEOR integrates time series RNA-seq and protein-DNA data to model temporal dynamics between TFs in an accessible and flexible framework for user exploration and to enable targeted follow-up biological experiments.

## Acknowledgements

We thank the National Science Foundation Graduate Research Fellowship Program and the Center for Computational and Molecular Biology for A.M.C. funding support. We thank the DRSC/TRiP Functional Genomics Resources, Harvard Medical School for hosting TIMEOR. We thank the Larschan, Lawrence, and Perrimon lab members for their advice. We thank the Adelman-Bender-Cole-Kuroda-Larschan joint lab meeting group for providing insights during presentations about this work. We thank Megan Gura and Dr. Anastasia Murthy for contributed ideas and manuscript comments. This research was supported by NIH grant to E.L.: 5R35GM126994.

## Author Contributions

A.M.C led the project, created back and front-end methods, created simulation, gathered data, led the data analysis, generated all tables, figures, and wrote manuscript. N.G. assisted in building part of front-end methods and wrote section of supplement. Y.H. provided informatics support and ideas. N.P. provided informatics support, the main data used in manuscript, and ideas. R.S. advised A.M.C. on the simulation. C.L. advised A.M.C. on modeling the data, methods development, and manuscript writing. E.L. advised A.M.C. on modeling the data, methods development, interpreting the biological mechanisms, and manuscript writing.

## Declaration of Interests

The authors declare no competing interests at the end of the manuscript.

## Lead Contact And Materials Availability

Further information and requests for resources and reagents should be directed to and will be fulfilled by the Lead Contact, Erica Larschan.

## TABLE OF CONTENTS

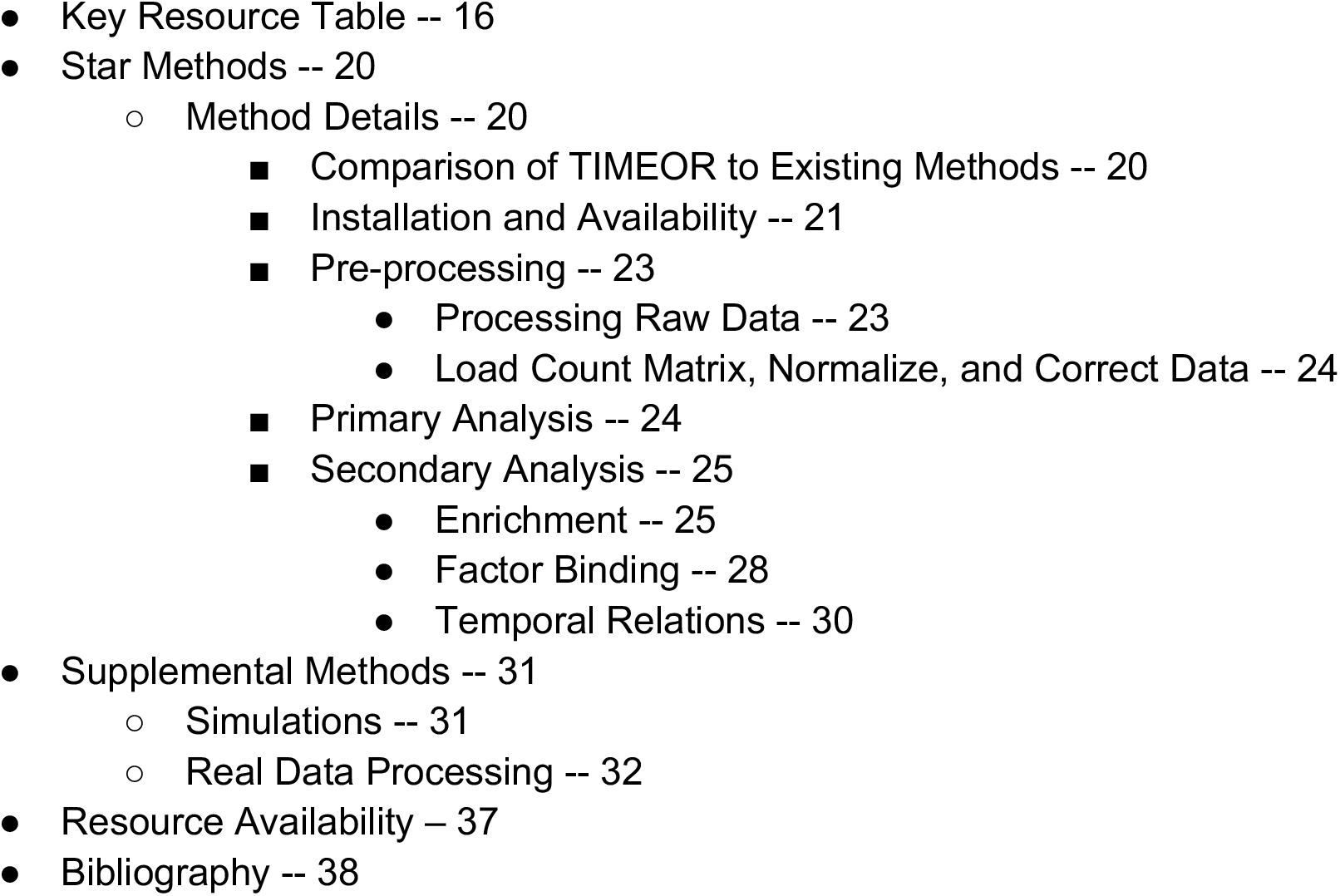

### KEY RESOURCE TABLE

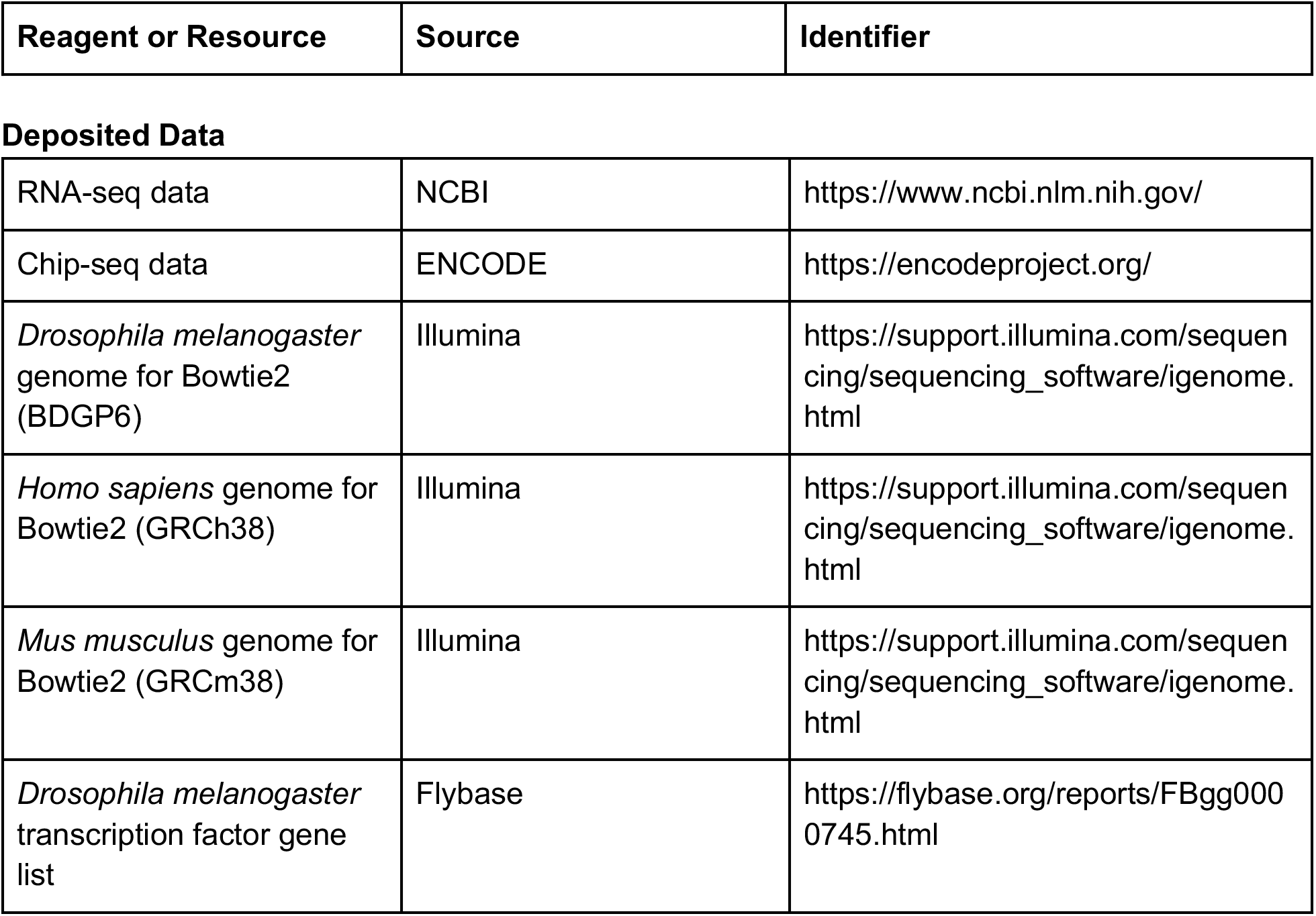
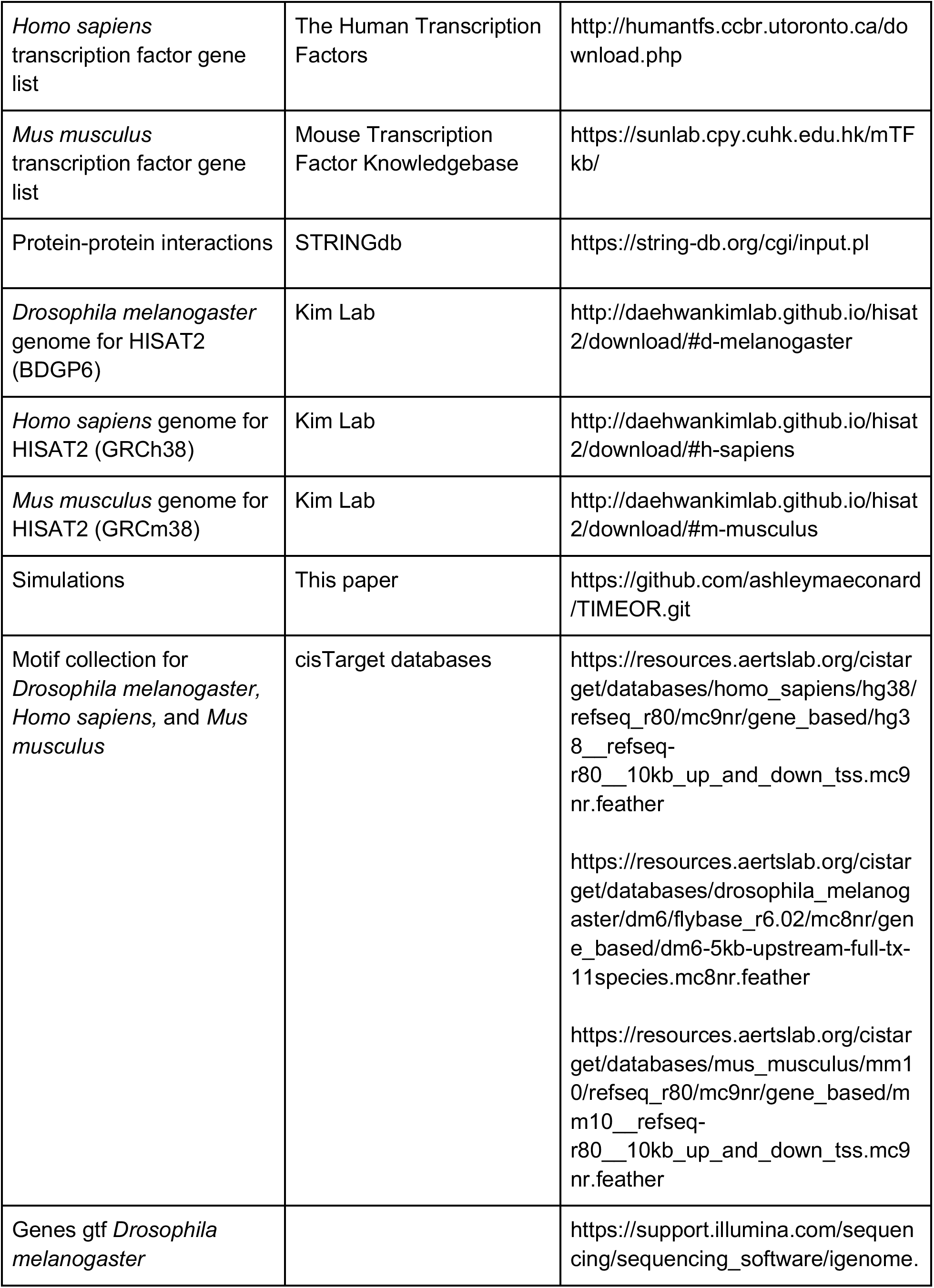
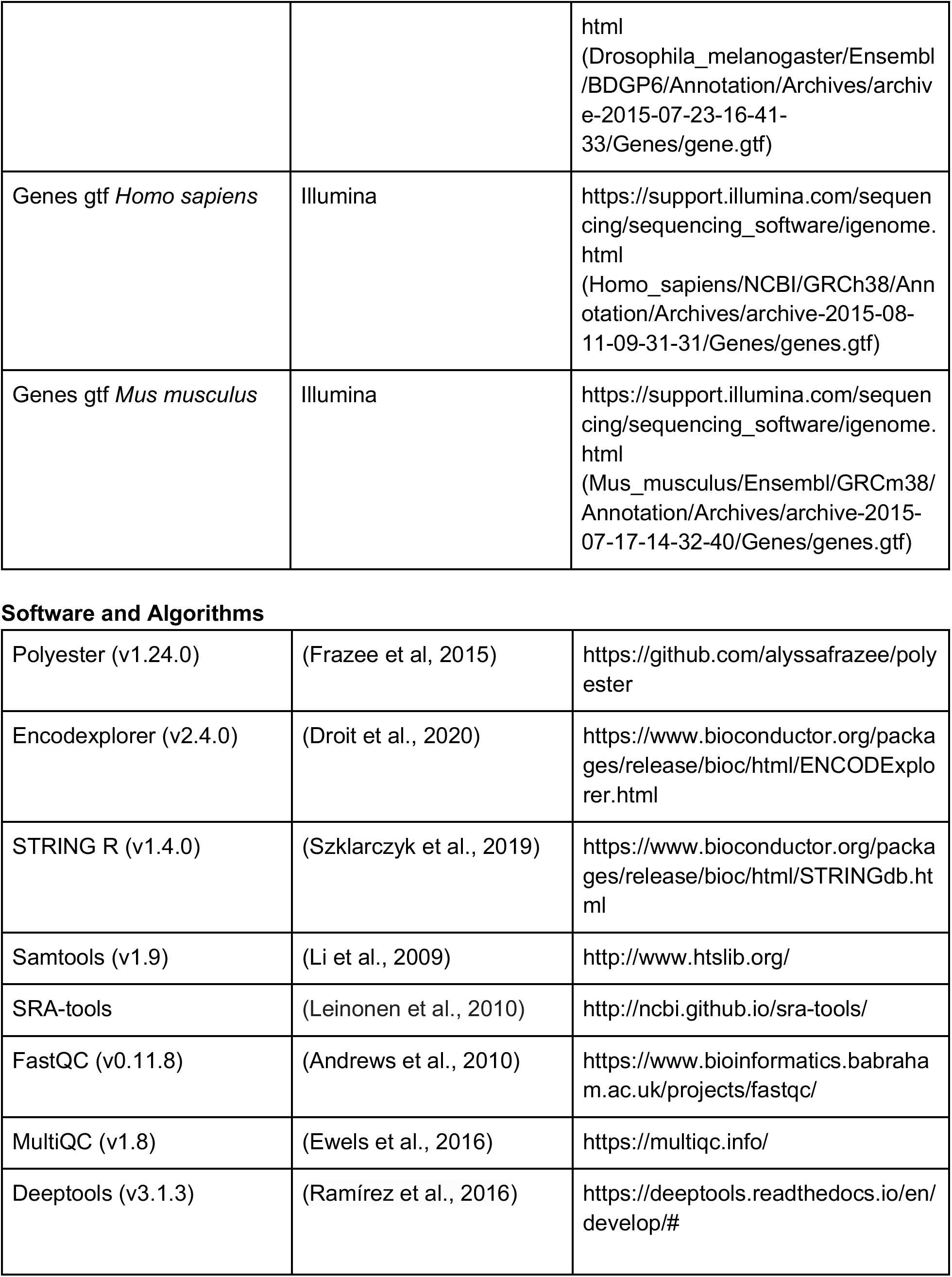
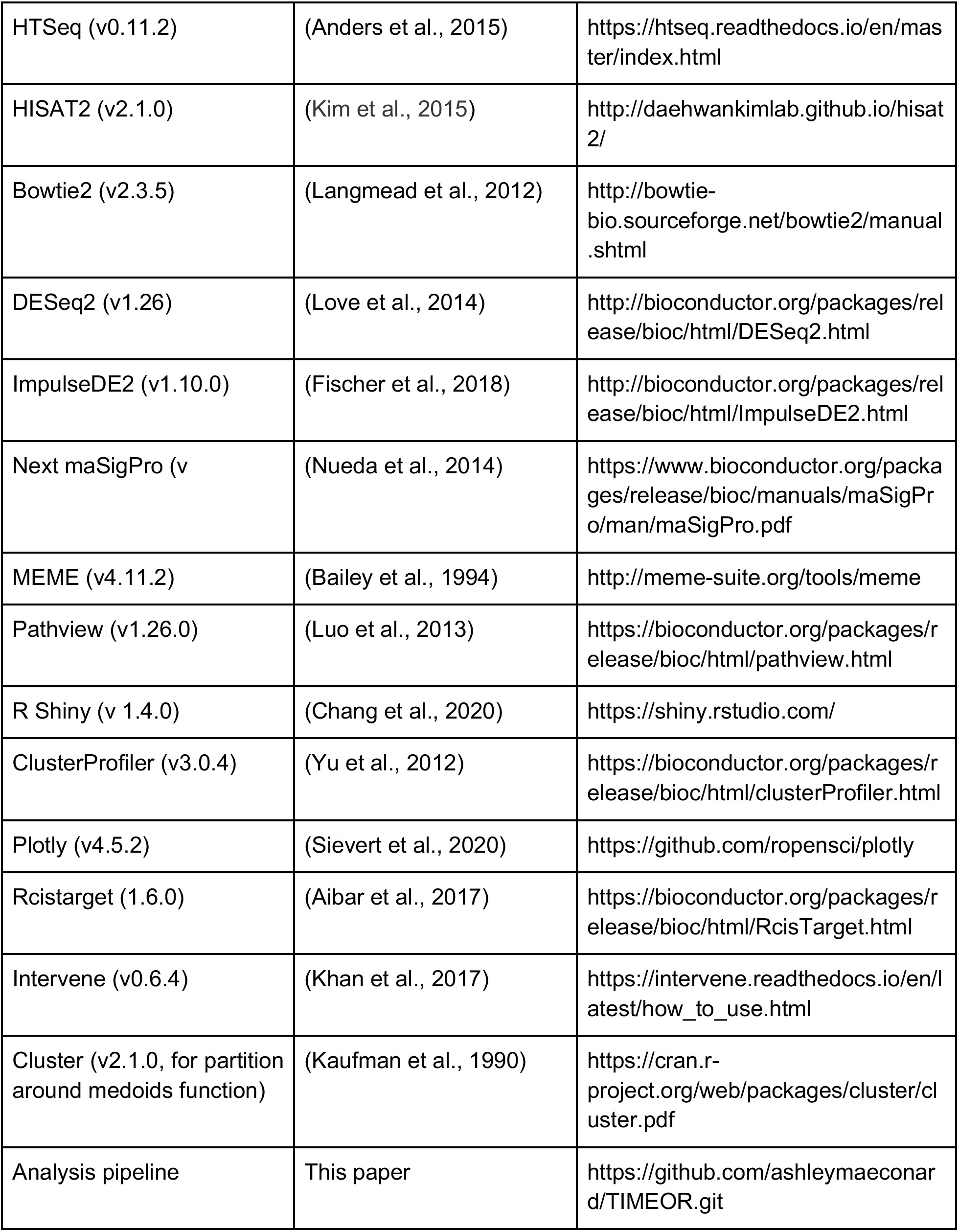

### STAR ✶ METHODS

#### METHOD DETAILS

##### COMPARISON OF TIMEOR TO EXISTING METHODS

This is the first pipeline method to focus on comprehensive and comparative analysis of time series and multi-omics data with the goal to infer gene regulatory mechanisms. **Figure S1** shows a comparison of multiple pipeline methods most similar to TIMEOR, including BioJupies and iDEP. We compared these methods in terms of 14 features (**Figure S1**). Some of these methods analyze RNA-seq data from raw .fastq files and perform gene ontology, claim to be able to analyze time series RNA-seq data, and/or report gene regulatory networks (Afgan et al., 2018; Torre et al., 2018; Ge et al., 2018; Cornwell et al., 2018; de Jong et al., 2015; Jensen et al., 2017; Kartashov and Barski, 2015; Spurney et al., 2020). TIMEOR fills a gap by providing at least twice as many features to uncover gene regulatory mechanisms from time series data compared to existing methods.

**Figure S1.**
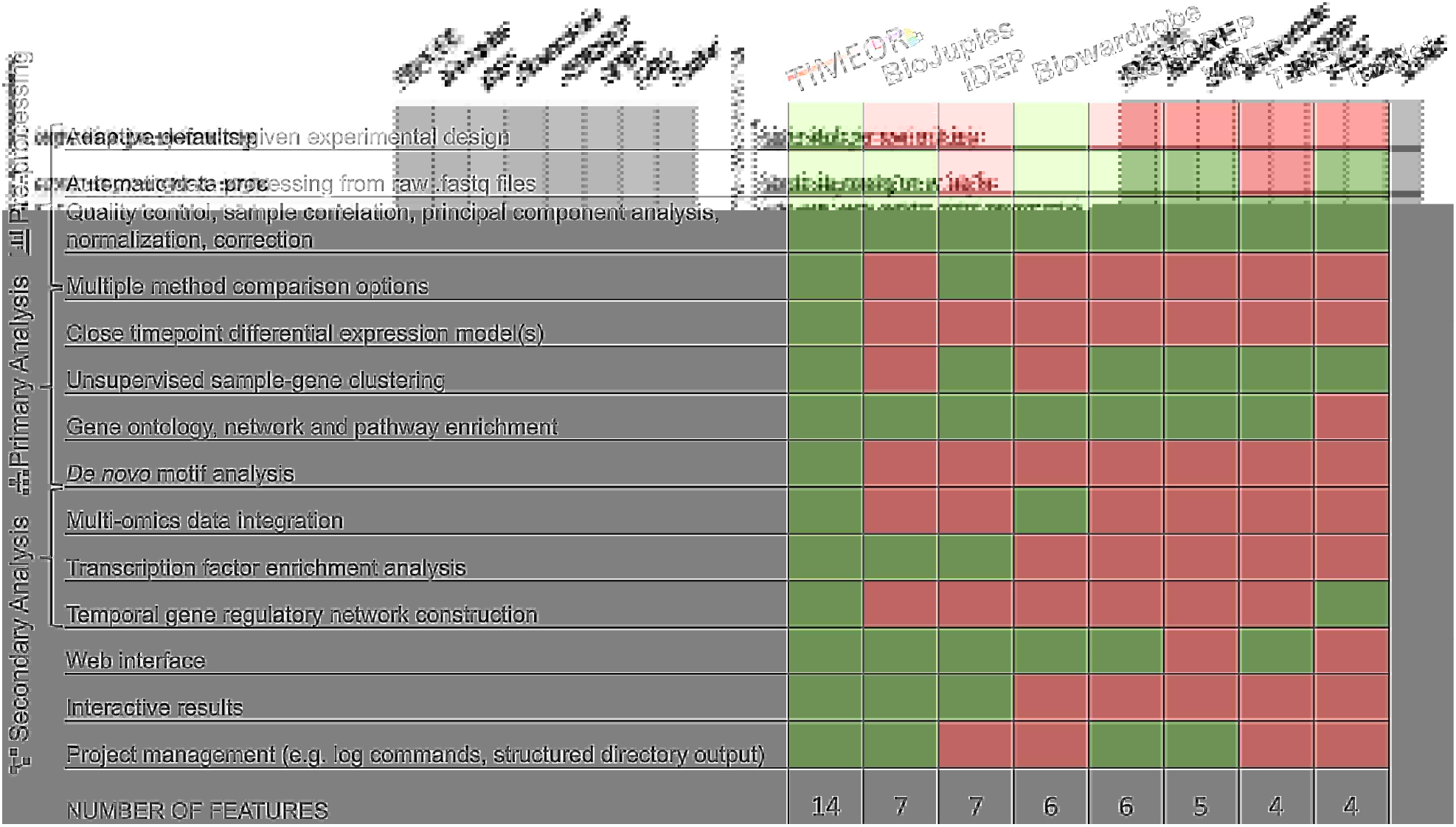
TIMEOR provides new functionalities to uncover regulatory network characteristics. Comparison between most similar pipeline methods where each feature (on left), or a close variant, is available (green) or not (red). The features are listed in a progressive fashion following the flow of data analysis.

As indicated in Figure S1, TIMEOR supports all aforementioned features. Specifically, TIMEOR pre-processes raw time series RNA-seq data by catering to its unique features through adaptive default methods before performing quality control and downstream processing (**Figure S2A, S2C, S2D**). This novel method also compares multiple methods for alignment and differential expression (DE, both categorical and continuous time points) to dynamically determine the best tools for the user’s analyses (**Figure S2B, S2E**). TIMEOR automatically determines the number of gene trajectory clusters before performing enrichment and *de novo* motif analyses on each cluster (**Figure S2E, S3A**). Because RNA-seq data alone does not provide information about direct interactions between transcription factors and their target genes, it is important to consider other types of omics data. To that end, TIMEOR predicts, provides and integrates transcription factor (TF) binding (mostly ChIP-seq) data from ENCODE (Davis et al., 2018; Joly Beauparlant et al., 2020) to help the user validate temporal gene-gene interactions (**Figure S3B**). Using these validated interactions, TIMEOR constructs the temporal TF gene regulatory network (**Figure S3D**). Overall, TIMEOR’s accessible RShiny (Chang et al., 2020) web interface enables researchers to determine the most suitable tools for their analyses, while producing paper ready and interactive figures. Furthermore, TIMEOR enables the user to inspect details on the methods used, their parameters and output, thereby facilitating reproducible research. See **Supplementary Data 3** for a table of itemized main features.

##### INSTALLATION AND AVAILABILITY

TIMEOR can be run through a conda environment (timeor_conda_env.yml). The web-app is going to be hosted online via (www.timeor.org) at DRSC/TRiP (https://fgr.hms.harvard.edu/tools).

To run TIMEOR, there are 3 prerequisites: 1) TIMEOR is downloaded; 2) R >= 3.6.1 is installed; 3) Anaconda/miniconda version 4.8.3 is installed.

1. Assuming Anaconda is installed in <CONDA_DIRECTORY>, in the command line type:

a. source <CONDA_DIRECTORY>/bin/activate
b. conda env create -f timeor_conda_env.yml
c. conda activate timeor_conda_env
2. Navigate to the TIMEOR folder and start R. Type:

a. library(shiny)
b. runApp(“app/”, launch.browser = F)
3. Proceed to the indicated address in your browser (preferably Chrome) to launch TIMEOR.

**Figure S2.**
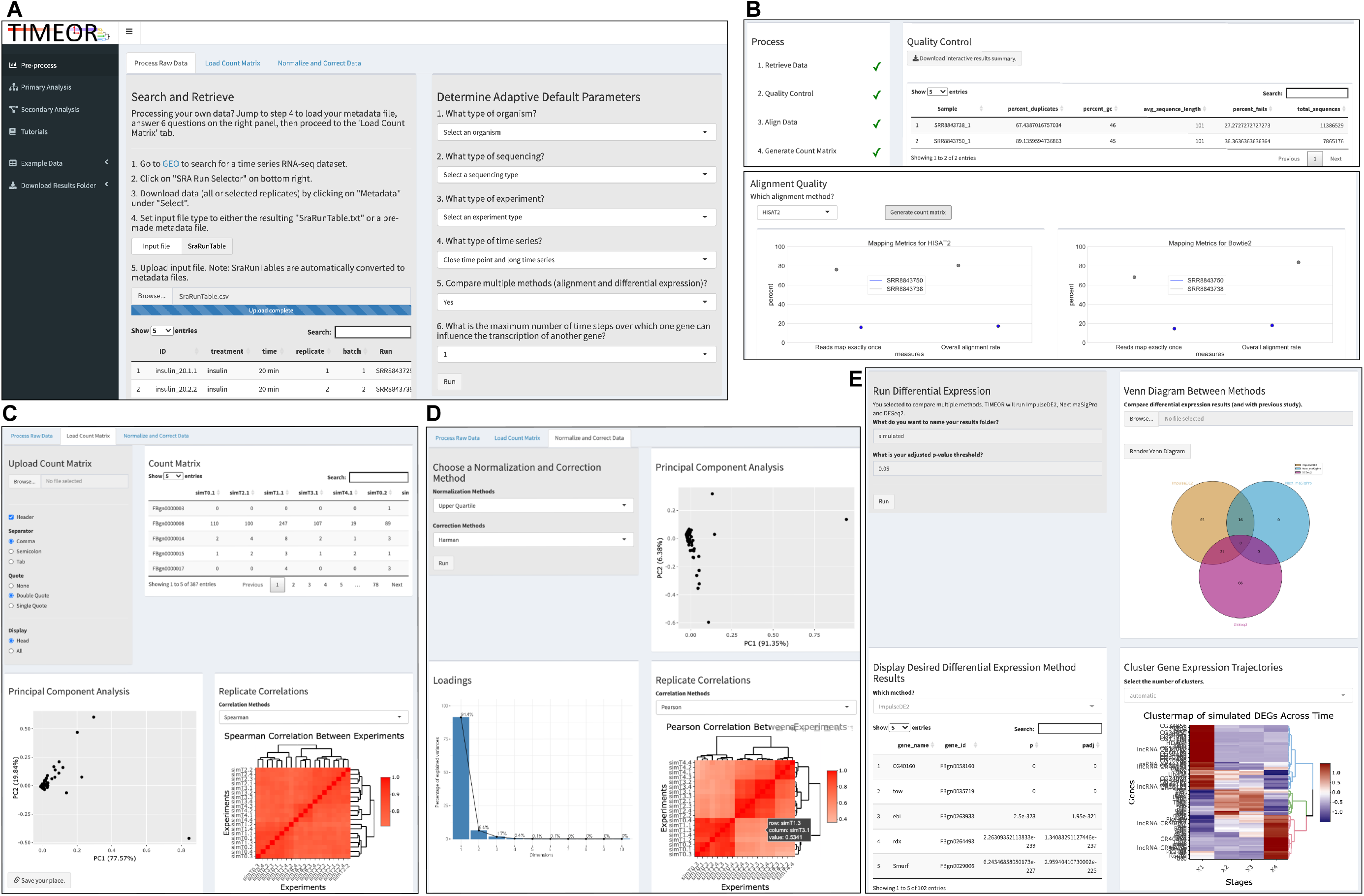
TIMEOR Application Pre-processing and Primary Analysis. **A.** In the Pre-processing stage in the “Process Raw Data” tab, the user chooses RNA-seq time series data and answers six questions about their data and desired analyses. TIMEOR subsequently sets the adaptive default methods that are customized to the user data. **B**. TIMEOR then retrieves and checks the data quality, providing interactive results to the user. These data are then aligned using either one or two alignment methods (Bowtie2 and HISAT2), and the user can choose which gives optimal results. The alignment .bam files are then used to create a gene-read count matrix which is then passed to the next tab “Load Count Matrix”. **C**. The gene-read count matrix is visualized through both interactive principal component analysis (PCA) and replicate (i.e. sample) correlation plots. Note that the user can pause the analysis at any time by clicking the “save your place” button and saving the link. **D.** The gene-read count matrix can then be passed to the “Normalize and Correct Data” tab where it is normalized (trimmed mean of M-values and upper quartile) and corrected. These altered data are again visualized using PCA and replicate (i.e. sample) correlation plots. **E.** The gene-read count matrix is passed to Primary Analysis (one tab only) where differential expression (DE) is performed using at least one of DESeq2, ImpulseDE2, and Next maSigPro, depending on how the user answered the adaptive default questions. The DE results between methods can be visualized using a Venn diagram. The user can also compare results with an outside gene list (e.g. from a past study) (**Figure 1B**). The results of each DE method are presented in the bottom left. The user can toggle to each method and TIMEOR will generate the associated gene trajectory clustermap. TIMEOR automatically chooses the number of clusters in the DE results. The number can also be chosen manually. Once the user chooses which DE method’s result to use, results are passed to Secondary Analysis (**Figure S3**).

##### PRE-PROCESSING – PROCESS RAW DATA

In the Pre-process Stage on the first tab “Process Raw Data”, the user is directed to the Gene Expression Omnibus (GEO) to find their desired time series RNA-seq data (**Figure S2A**). The user is instructed to upload the GEO generated data information file (SraRunTable.txt), which TIMEOR parses into a metadata file and generates the user’s personal analysis session folder for use in subsequent analysis. The user is instructed to answer six questions about their data. Namely, 1) “what type of organism (*Drosophila melanogaster, Homo sapiens*, or *Mus musculus)”*, 2) “what type of sequencing (paired-end or single-end)”, 3) “what type of experiment (case vs. control, or just case or control)”, and 4) “what type of time series (close time point and long time series, distant time point, and close time point and short time series)”. In addition, TIMEOR can compare two methods for multiple alignment (HISAT2 (Kim et al., 2015) and Bowtie2 (Langmead et al., 2012)) (**Figure 1A, S2B, S4A, S4B**) and several methods for differential expression (DE, DESeq2 (Love et al., 2014), ImpulseDE2 (Fischer et al., 2018), and Next maSigPro (Nueda et al., 2014) (**Figure 1B, S2E, S4C**), should the user choose ‘yes’ to 5) “compare multiple methods (that is, both alignment and DE methods)”. Lastly TIMEOR infers direct interactions between genes up to the maximum number of time steps input by the user by asking: 6) “What is the maximum number of time steps over which one gene can influence the transcription of another gene?”.

Once the user clicks “Run”, TIMEOR begins processing the answers to the six questions to choose the most appropriate adaptive default methods to run, followed by retrieving the raw .fastq data supplied in the uploaded GEO file. The retrieved .fastq files are then run through quality control using FastQC (Andrews et al., 2010) and summarized using MultiQC (Ewels et al., 2016). TIMEOR outputs the results in a summary table and interactive results for download. Next, each .fastq file (i.e. sample) is aligned to the reference genome for the organism designated in question 1. If the user chose to compare alignment methods, an alignment plot is output to the screen showing the overall alignment scores for each sample for both alignment methods (**Figure S2B**). If the user chose not to compare, HISAT2 (Kim et al., 2015) alignment results are plotted to show the uniquely mapped read percentage and overall alignment. These alignment results are then passed to HTSeq (Anders et al., 2015) to produce a read count matrix of samples by read counts per gene. Then for each sample and for each gene, a read count method is used to calculate the number of mapped reads to each gene, resulting in a vector of transcript counts per gene for each sample. All read count vectors for all samples are then merged to form one large transcript count matrix of genes by samples. TIMEOR indicates that each of the aforementioned steps is complete with a check mark (**Figure S2B**).

##### PRE-PROCESSING – LOAD COUNT MATRIX, NORMALIZE AND CORRECT DATA

The read count matrix is then passed to the next tab “Load Count Matrix” (**Figure S2C**) where filtering and principal component analysis (PCA) are performed. Importantly, the user can also begin on this tab by simply loading the pre-computed read count matrix and associated metadata file. TIMEOR visualises PCA in an interactive plot, along with a loadings plot below it. TIMEOR performs replicate (or sample) correlations using either the Spearman or Pearson correlation, which are displayed in an interactive symmetrical heatmap (**Figure S2C**).

This read count matrix can be normalized and corrected on the next tab “Normalize and Correct Data” (**Figure S2D**). The user can choose between two normalization methods: trimmed mean of M-values, and upper quartile, and correction is performed by Harman (Oytam et al., 2016). Once the user clicks “Run”, TIMEOR outputs the resulting PCA plots of the normalized and corrected read count data. The user can again perform replicate (or sample) correlations on the normalized and corrected data. Note that at any point in this analysis, the user can save their current location by clicking on the button at the bottom on each web-page (**Figure S2C** bottom left).

##### PRIMARY ANALYSIS

In the Primary Analysis stage, the user is asked to name the results folder and set up the adjusted p-value threshold (**Figure S2E**). If the user selected to compare multiple differential expression methods (as part of the six initial questions to set the adaptive default methods) and the data are close time series, TIMEOR will run DESeq2 (Love et al., 2014), ImpulseDE2 (Fischer et al., 2018), and Next maSigPro (Nueda et al., 2014). If the user selected that these timepoints are distant, DESeq2 (Love et al., 2014) will be run using the gene-by-sample read count matrix. Note that TIMEOR inputs the raw read count matrix for DESeq2 (Love et al., 2014) and ImpulseDE2 (Fischer et al., 2018), as they require non-normalized and corrected read count data and the pre-normalized and corrected data for Next maSigPro (Nueda et al., 2014). When the user clicks “Go” the method(s) are run. Once finished, the user can compare the overlap of DE genes between methods in a Venn diagram (**Figure S2E**). The user can also compare these results with a list of genes (from a past study for example). TIMEOR outputs the results for each method in a table that can be selected using the dropdown menu. When a method is selected, TIMEOR produces the associated gene trajectory clustermap on the right (**Figure S2E**).

The clustermap (Sievert et al., 2020) interactively highlights clusters of DE genes’ expression trajectories in the form of a log2 fold change or z-score, formed using Euclidean distance and Ward.D2 (Murtagh et al., 2014) hierarchical clustering. Importantly, TIMEOR addresses the challenge of clustering by taking the mode of 3 unsupervised clustering methods (partition around medoids (Reynolds et al., 2006), Silhouette (Rousseeuw et al, 1987), and Calinski criterion (Caliński et al., 1974)) to automatically return the number of gene trajectory clusters to the user. Given that clustering is an NP-hard problem (Křivánek et al., 1986), TIMEOR also provides an Elbow plot (**Figure S4D**) to aid the user in choosing the number of clusters manually. Once the user determines which DE method to use, those results are passed to Secondary Analysis to determine temporal relations between genes and TFs.

##### SECONDARY ANALYSIS

TIMEOR performs three types of Secondary Analysis, all in separate tabs. Namely “Enrichment” (**Figure S3A**), “Factor Binding” (**Figure S3B**), and “Temporal Relations” (**Figure S3D**) to uncover both what processes are enriched (i.e. affected) in the user’s time series experiment, and how those genes are regulating that enriched process. In the following sections we describe the details of each tab separately.

##### SECONDARY ANALYSIS – ENRICHMENT

**Figure S3:**
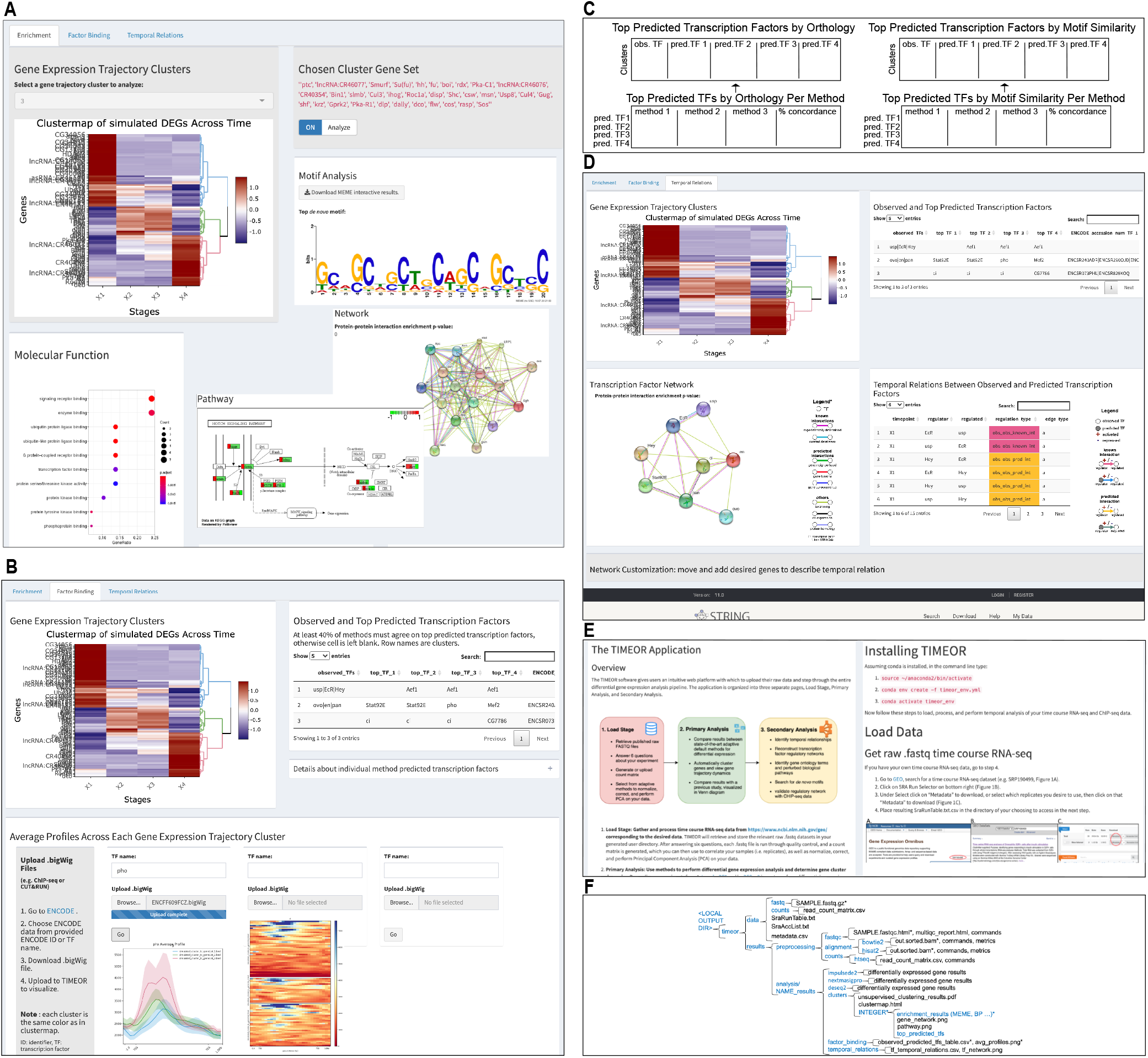
TIMEOR Application Secondary Analysis. **A**. In the Secondary Analysis Stage “Enrichment” tab, each gene trajectory is assessed for gene ontology, pathway, network, and de novo motif enrichment. **B.** TIMEOR scans for the observed and top four predicted transcription factors (TFs) to bind to each gene trajectory cluster (table top right) from the *Top Predicted Transcription Factor Binding by Orthology* table. TIMEOR returns ENCODE IDs where ChIP-seq data exist for each TF. The user can then choose which TFs to plot over each gene trajectory cluster to help validate key regulators. **C.** TIMEOR displays four predicted TF tables: 1) *Top Predicted Transcription Factors by Orthology* table, giving information regarding: the observed, predicted TFs by orthology, and associated ENCODE IDs (not shown), which is built from a consensus between multiple methods’ results, visible in 2) *Top Predicted Transcription Factors by Orthology per Method* table consisting of the top predicted TFs for each individual method by orthology. If there is a consensus among methods in table 2 for any of the top four TFs above 40%, those results are displayed in table 1. 3) *Top Predicted Transcription Factors by Motif Similarity* table, giving information regarding: the observed, predicted TFs by motif similarity, and associated ENCODE IDs (not shown), which is built from a consensus between multiple methods’ results, visible in 4) *Top Predicted Transcription Factors by Motif Similarity per Method* table, consisting of the top predicted TFs for each individual method. If there is a consensus among methods in table 3 for any of the top four TFs above 40%, those results are displayed in table 4. **D.** TIMEOR infers the temporal relations between the observed and predicted transcription factors. The user can move nodes in the desired order using the STRINGdb interface. **E.** TIMEOR provides two tutorials, one for the web-app and one for the command line version of TIMEOR. **F.** Each session’s results are clearly organized in the user’s personal analysis session folder to download for future use. Blue indicates a folder; black indicates a file, webpage, or image; and an asterisk indicates many files or folders of the same type.

The resulting gene trajectory clusters are passed to the “Enrichment” tab in the “Secondary Analysis” stage. The user can toggle through the clusters to see the gene trajectory cluster (i.e. gene set) they wish to analyze (**Figure S3A**). Once the user clicks on the analyze button, the following enrichment analyses are performed: gene ontology (GO), pathway, network, and *de novo* motif. For GO analysis, ClusterProfiler (Yu et al., 2012) is run with a Benjamini Hochberg adjusted p-value cutoff of 0.05. If particular GO terms (molecular function, biological process, and cellular component) are significantly enriched for that group of genes, those results are depicted as “dot plots” under the clustermap on the “Enrichment” tab. *Dot plots* highlight the enriched GO terms (x-axis) vs. the ratio of the genes within the cluster that are enriched for that GO term (y-axis). The radius of the dot shows the count of genes enriched for that GO term and the color of the dot indicates how significant the GO term is.

TIMEOR stores other GO result formats including *ontology plots*, where the leaves of the tree are the most specific enriched terms, and a *relationship graph*, highlighting which GO terms are related to each other. The radius of each node shows the count of genes enriched for that GO term and the color of the dot indicates how significant the GO term is. TIMEOR also outputs spreadsheets of all results into the user’s personal analysis session folder.

Next, pathway level analysis is performed on each selected gene trajectory cluster using PathView (Luo et al., 2013), where an enriched pathway (Benjamini Hochberg adjusted p-value ≤ 0.05) is output with temporal highlighting for genes (−1 is most downregulated and 1 is most upregulated).

TIMEOR then uses STRINGdb (Franceschini et al., 2013) to search for enriched protein-protein interaction networks, if the enrichment for that group of genes is below a q-value ≤ 0.05. STRINGdb highlights various types of known interactions between genes, namely ‘experimentally determined’ and from ‘curated databases’. StringDB also highlights predicted edges from ‘gene neighborhoods’, ‘gene fusion’, and ‘gene co-occurrence events’. Lastly STRINGdb provides ‘text-mining’, ‘co-expression’, and ‘protein-homology’ interaction edge types. TIMEOR also stores a table of these interactions, along with the PubMed IDs that support ‘experimentally determined’ edges in the user’s personal analysis session folder.

Lastly, TIMEOR uses MEME (Bailey et al., 1994) to discover novel (i.e. *de novo)*, ungapped motifs in the sequences within each gene trajectory cluster, and returns the top three motifs. The positive strand motifs are shown, along with MEME’s downloadable interactive results. TIMEOR stores the reverse complement motifs, and each gene’s DNA sequence files in the user’s personal analysis session folder. If the user desires, TIMEOR’s command line tools can be run to scan each cluster to output any RNA types which are significantly enriched within each gene cluster using the hypergeometric test of significance. The RNA types are lncRNA, miRNA, pre-miRNA, snRNA, snoRNA, tRNA, pseudogene, rRNA, and protein coding.

##### SECONDARY ANALYSIS – FACTOR BINDING

On the next tab “Factor Binding” within Secondary Analysis, TIMEOR scans across all clusters to identify both observed and predicted TFs (**Figure S3B**), and list any ENCODE (Davis et al., 2018; Joly Beauparlant et al., 2020) IDs associated with predicted TF binding data. TIMEOR displays the following four predicted TF tables: 1) *Top Predicted Transcription Factors by Orthology* table, consisting of three pieces of information: a. observed TFs, predicted TFs by orthology, and c. associated ENCODE IDs. 2) *Top Predicted Transcription Factors by Orthology per Method* table lists the four top predicted TFs by orthology for each individual method. 3) *Top Predicted Transcription Factors by Motif Similarity* table lists three pieces of information: a. observed TFs, b. predicted TFs by motif similarity, and c. associated ENCODE IDs. 4) *Top Predicted Transcription Factors by Motif Similarity per Method* table lists the four top predicted TFs by motif similarity for each individual method (**Figure S3C**). In what follows we describe how each table is made.

TIMEOR performs this TF prediction using RcisTarget (Aibar et al., 2017) to first identify TF binding motifs which are over-represented (i.e. enriched) in a gene list. RcisTarget uses a database (**Key Resource Table Deposited Data**) containing genome-wide motif rankings, collected through multiple methods including HOCOMOCO (Kulakocskiy et al., 2017), Transfac (Wingender et al., 1996), Jaspar (Sandelin et al., 2004), CIS-BP (Weirauch et al., 2014), HOMER (Heinz et al., 2010), and several labs’ datasets including Dr. Jussi Taipale (Aibar et al., 2017). RcisTarget performs motif-enrichment analysis on a gene list by first estimating the over-representation of each motif on the gene set. RcisTarget displays the resulting selection of significant motifs in ranked order by the Normalized Enrichment Score (NES). This is calculated for each motif based on the area under the curve distribution of all the motifs for the gene-set. Those motifs that pass the given threshold (3.0 by default by changeable by user) are considered significant.

Rcistarget’s results are mostly long lists of redundant significantly enriched motifs and associated candidate TFs that are derived from multiple databases. TIMEOR prioritizes this long list by first forming a consensus about the top four (default but changeable by user) candidate TFs which bind to the input group of genes. This consensus is formed in three steps. First, enriched motif search is a widely studied problem, and TIMEOR leverages this fact to scan across all motif databases for which TF binding motifs are enriched. TIMEOR does this by first splitting the RcisTarget ranked list results into smaller ranked lists, one for each database used to identify enriched motifs. If at least 40% (default but changeable by user in command line version of TIMEOR) of the databases agree on their top ranked TF, then ENCODExplorer (Joly Beauparlant et al., 2020) is used to scan for any ChIP-seq data that exists for this TF. This procedure continues for the second, third, and fourth candidate TFs. TIMEOR repeats the run of RcisTarget and ENCODExplorer on each cluster to identify its recommendation for at most the top four candidate TFs. This process is repeated to find two types of candidate TFs.

Each enriched motif is associated with candidate TFs of two kinds, and are output into two tables: 1) based on orthologous sequences and placed in *Top Predicted Transcription Factors by Orthology* table, and 2) based on similarities between annotated and unknown motifs and placed in *Top Predicted Transcription Factors by Motif Similarity* table (Aibar et al., 2017; Verfaillie et al., 2014). TIMEOR then uses ENCODExplorer to find associated factor binding data from ENCODE where possible. The user can also see the associated *Top Predicted Transcription Factors for Method* table, consisting of the top predicted TFs for each individual method for both orthology and motif similarity (**Figure S3C**).

**Figure S4:**
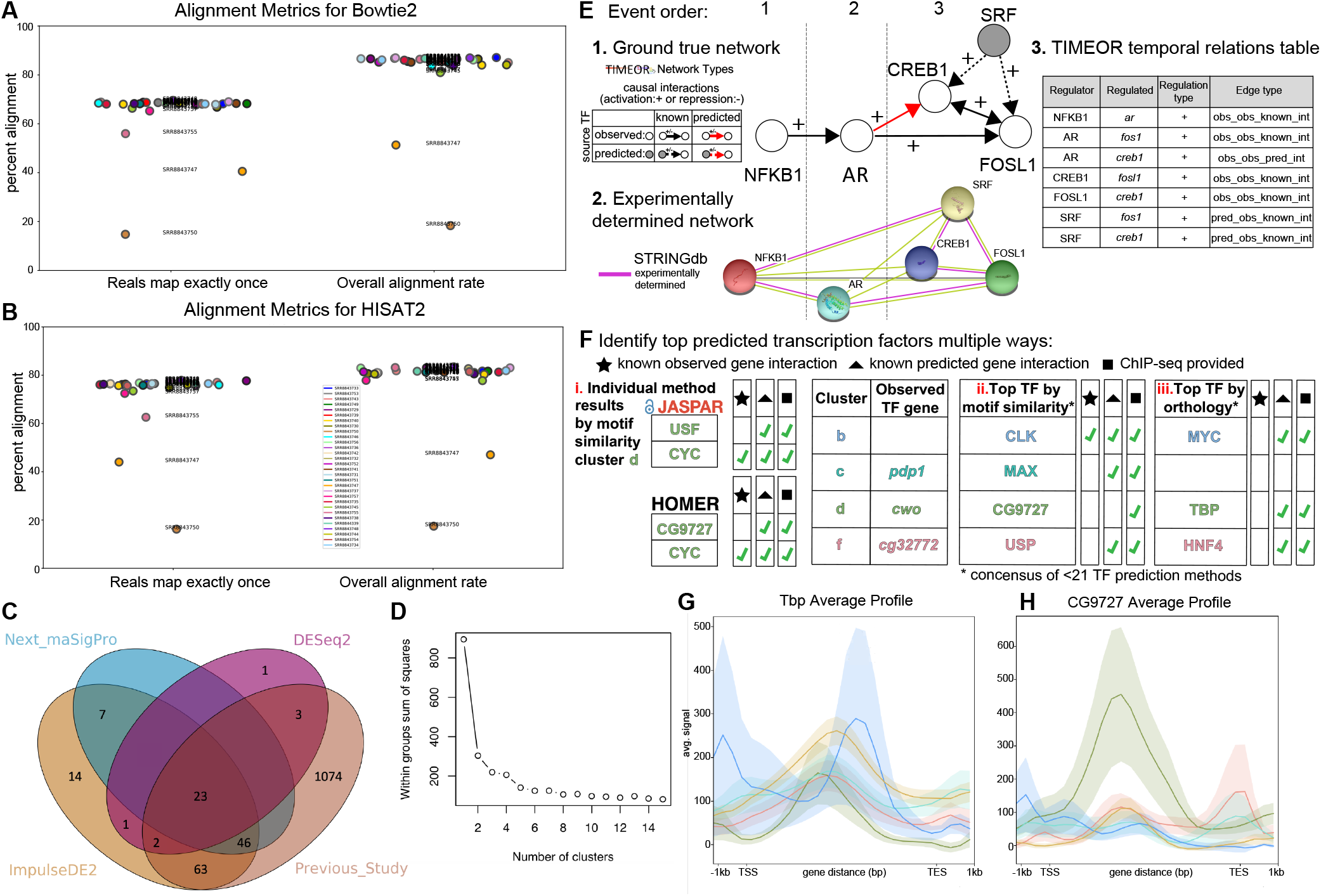
TIMEOR’s visual outputs assist the user to make analysis decisions. **A**. Alignment metrics from HISAT2 for data from Zirin et al. 2019. **B**. Alignment metrics from Bowtie2 for data from Zirin et al. 2019. **C**. TIMEOR produces a Venn diagram comparing categorical (DESeq2) and continuous (ImpulseDE2, Next maSigPro) differential expression methods and the previous study (Zirin). ImpulseDE2 shows the highest percent overlap with the previous study and other methods. **D**. TIMEOR uses multiple methods to assess stability of the number of clusters. The Elbow method showed decreasing error stabilized at six clusters. **E**. Simulation design, 1. The simulated ground truth network, 2. STRINGdb’s TF network highlighting *known* (i.e. experimentally determined) interactions, and 3. TIMEOR’s ground truth temporal relations table (gene regulatory network). **F**. Multiple predicted TF tables were used to help identify the top predicted TFs binding to each cluster from Zirin et al., 2019 data. In i. are the top transcription factors called by popular TF prediction methods highlighting that CYC has known observed and predicted TF interactions and ChIP-seq data to validate interactions. In ii. are the top transcription factors by orthology (table titled *Top Predicted Transcription Factors by Orthology)*, and in iii. are the top transcription factors by motif similarity (table titled *Top Predicted Transcription Factors by Motif Similarity)*. Each table also provides any ENCODE identifiers for ChIP-seq data if applicable (Davis et al., 2018; Joly Beauparlant et al., 2020). **G**. TBP average ChIP-seq profile over the gene body ±1KB for all six clusters. **H**. CG9727 average ChIP-seq profile over the gene body ±1KB for all six clusters.

TIMEOR then prompts the user to upload their own or processed (using the ENCODE IDs provided by TIMEOR) TF binding data in the form of a .bigWig file, to see how the candidate TF(s) bind to genes within each gene trajectory cluster. DeepTools (Li et al., 2009) is used to generate an average profile and heatmap over the gene body ±1kb from the transcription start and end site (**Figure S3B**). TIMEOR can process three TFs at a time through the web interface. All results are stored in the user’s personal analysis session folder.

##### SECONDARY ANALYSIS – TEMPORAL RELATIONS

On the “Temporal Relations” tab, TIMEOR harnesses the time series RNA-seq data which TIMEOR clustered, STRINGdb’s “experimentally determined” edges, and knowledge of the observed and predicted TFs that bind each cluster. Specifically, the observed and predicted TFs within the *Top Predicted Transcription Factors by Orthology* table are combined with the temporal dynamics across gene trajectory clusters to form the “Temporal Relations Between Observed and Top Predicted Factors” table. This latter table defines the GRN and specifically highlights four relationships in the form of a quadruple (source TF, target TF, regulation type, interaction type). *Interaction types* are: “predicted to observed known interaction”, “predicted to observed predicted interaction”, “observed to observed known interaction”, or “observed to observed predicted interaction”. *Regulation types* are “activation” or “repression”. Specifically, if a predicted TF interacts with an observed TF and this is a known interaction (as specified by STRINGdb’s experimentally determined edges) then that gene is denoted to have an edge type “pred_obs_known_int” (i.e. predicted TF and observed TF have known interaction). Furthermore, this edge can show that the predicted TF is activating or repressing the observed TF. Similarly, an observed TF can interact with another observed TF through a known interaction (denoted “obs_obs_known_int”). TIMEOR proposes possible novel interactions between a observed TF activating or repressing another observed TF, or a predicted TF activating or repressing a observed TF. These edge types are denoted “obs_obs_pred_int” and “pred_obs_pred_int”, respectively (**Figure S4D**).

Note that TIMEOR only evaluates the regulation type of observed TFs. Importantly, these interactions are confined to a window of time set by the user for question six: “What is the maximum number of time steps over which one gene can influence the transcription of another gene?” to determine the window of time in which to look for potential causal relations between TFs. Measuring such causal relations is only truly feasible using time series RNA-seq data. TIMEOR’s simple algorithm takes the time window defined by the user in question six, and reports which TFs might influence each other, and how in that window. More specifically, TIMEOR reports which interactions are *known* between observed TFs and observed and predicted TFs, and *predicts novel* interactions between observed TFs, while also indicating each repressive or activating effect. TIMEOR’s flexible framework enables the user to change this window threshold as desired.

At the bottom of this last tab TIMEOR provides the STRINGdb web-interface to facilitate the user to visualize and customize the resulting temporal regulatory network. In this way, the user can add and move TF nodes around, as well as add other non-TF genes to the network (**Figures 3D, S3D**).

### SUPPLEMENTAL METHODS

#### SIMULATIONS

Polyester version 1.24.0 (Frazee et al., 2015) was used to simulate four RNA-seq expression cascading activation patterns over six time points (**Figure S3E**) two biological replicates for 63 genes at approximately 20X sequencing coverage. Note that the first time point was used as the control. Those fold change expression cascades were trajectory 1: 0.5,2,2,3,4,4.5; 2: 0.2,0.5,2,4,4.5,5; 3: 0.1,0.5,0.8,0.9,1,3; and 4: 0.05,0.1,0.1,0.5,0.5,4. To obtain an accurate baseline gene expression across the five timepoints, we analyzed a five time point RNA-seq experiment of fusobacterium nucleatum-stimulated human gingival fibroblasts control samples (GEO: GSE118691) taken from two donors (1 and 2) at 2, 6, 12, 24, 48 hours (Kang et al., 2019a; Kang et al., 2019b). We simulated a constant RNA-seq expression of one fold across all five timepoints for all those genes that were active in at least one time point. This culminated in simulating expression for 17723 genes of which four are observed TFs and one TF is predicted. That is, the fourth expression trajectory activated last was enriched for the predicted TF SRF to bind by 22 genes (Han et al., 2018).

To simulate the temporal relationships between these five TFs, TIMEOR enables the user to choose the window of possible interaction between TFs along the time course, which we set to be one time point. In this window, Figure S3E step 1 highlights our ground truth simulation of four observed TFs (one for each activation cascade trajectory) and one predicted TF. Specifically, at the first time point, we simulated initial activation of AR by NFKB1 which is a *known* (i.e. experimentally determined) interaction (**Figure S3E**). At the second time point we simulated AR activating FOSL1 (known interaction) and CREB1 (predicted interaction). At the third time point, we simulated SRF to activate the two observed genes (FOSL1 and CREB1) in known interactions. At this same time point, we simulated FOSL1 and CREB1 to activate each other in known interactions.

This simulation can be described in nine steps. First, as input, Polyester takes annotated transcript nucleotide sequences in the form of cDNA sequences in FASTA format. We used bioMart (https://m.ensembl.org/biomart/martview/) to extract the first transcript sequence for each of 63 genes. Second, TIMEOR ran Bowtie2 version 2.3.5 (Langmead et al., 2012) to align all sample .fasta files to the human genome (GRCh38):

> (bowtie2 -f -p 3 -x /genomes_info/hsa/genome_bowtie2/genome -U sample_t01.fasta --un-gz out_un.sam.gz --al-gz out_al.sam.gz --met-file out_met-file.tsv -S out.sam 2> summaryfile.txt),

and achieved an average 83.9% alignment rate. Third, samtools version 1.9 (Li et al., 2009) then converted and sorted all .sam files to .bam files:

> (samtools view -S -b out.sam > out.bam; rm -rf out.sam out.sorted.sam; samtools sort out.bam -o out.sorted.bam; rm -rf out.bam).

Fourth, TIMEOR ran HTSeq to produce a read count matrix of samples by read counts per gene for each sample, and TIMEOR then merged the samples in chronological order to create the final read count matrix of genes by sample (i.e. replicates).

> (htseq-count -f bam -r pos -i gene_id out.sorted.bam BDGP6/genes.gtf > htseq_counts).

Fifth, TIMEOR ran ImpulseDE2 version 1.10.0 (Fischer et al., 2018) at an adjusted p-value threshold of 0.05 and found 65 DE genes. Neither of the additional genes were TFs and thus not selected for downstream TF temporal regulation analysis. Sixth, TIMEOR clustered the gene trajectories for all DE genes into 4 clusters. Seventh, for each cluster TIMEOR identified and summarized the top four (default by changeable by user) predicted TFs across at most 21 TF prediction methods when toggling the percent concordance at various thresholds (2, 5, 15, 25, 35, 45, 55, 65, 75, and 85 percent), and reported the percent concordance among these methods’ ranks for the top four predicted TFs. Note that TIMEOR uses only the methods that output a TF when calculating percent concordance. Eighth, TIMEOR infers the temporal relationships between observed and top one predicted TFs for each cluster. This process of calling the top predicted TFs and inferring the temporal relations happened 10 times by toggling this percent concordance parameter between 2% - 85%.

Ninth, we calculated recall and precision to assess TIMEOR’s robustness to recover the true underlying (i.e. ground truth) gene regulatory network (GRN) after determining the TF temporal relationships. We defined the quadruple (source TF, target TF, regulation type, interaction type). *Interaction types* are: “predicted to observed known interaction”, “predicted to observed predicted interaction”, “observed to observed known interaction”, or “observed to observed predicted interaction”. *Regulation types* are “activation” or “repression”. A true positive consists of all correct quadruples. A false negative is failing to report one of the ground truth quadruples. And a false positive consists of introducing a new quadruple. TIMEOR recovered the true network with perfect recall at all percent concordance thresholds except at high percent concordance (65% and above) which lead to just one ground truth TF to drop out. As expected, at low percent concordance between TF prediction methods TIMEOR predicted other TF interactions which were not simulated (i.e. not part of ground truth). At a concordance threshold of at least 35 and above we obtained perfect precision (**Figure 2A**).

#### REAL DATA PROCESSING

TIMEOR uncovered the complex molecular functions after insulin stimulation in *Drosophila* S2R+ cell (project ID SRP190499, **Supplementary Data 1**) by integrating both temporal RNA-seq and ChIP-seq data (**Supplementary Data 2**). Through TIMEOR, the steps described next were followed. The data were gathered from NCBI project SRP190499 using fastq-dump (Li et al., 2009). Note only 2 of 30 samples shown. The specific command used was:

> (parallel -j $1 --bar fastq-dump {1} -split-files -O $2 -gzip ::: SRR8843729 SRR8843730).

The sequenced reads were run through FastQC (Andrews et al., 2010) with default parameters to check the quality of raw sequence data and filter out any sequences flagged for poor quality. No sequences were flagged. The specific command used was:

> (fastqc SRR.fastq.gz --outdir=/fastQC_results/)

Sequenced reads were then mapped to release 6 *Drosophila melanogaster* genome (dm6) using Bowtie2 (Langmead et al., 2012) and HISAT2 (Kim et al., 2015) (**Figure S4A, B**). Bowtie2 was used to align reads to the dm6 genome with this command:

> (bowtie2 -p 3 -x /genomes_info/dme/genome_bowtie2/genome -U SRR.fastq.gz --un-gz out_un.sam.gz --al-gz out_al.sam.gz --met-file out_met-file.tsv -S out.sam 2> summaryfile.txt)

HISAT2 was used to align reads to the dm6 genome, followed by sorting and converting to a bam file:

> (hisat2 -p 3 --dta -x /genomes_info/dme/genome_hisat2/genome -U SRR.fastq.gz -t --un-gz out_unalign.sam.gz --al-gz out_al_atleastonce.sam.gz --known-splicesite-infile /genomes_info/dme/splicesites.txt --novel-splicesite-outfile novel_splicesite --summary-file summaryfile.txt --met-file met-file.txt -S out.sam 2> alignmentsummary.txt).

HISAT2 was chosen due to higher unique read mapping (**Figure S4A, S4B**). All sample .sam files were sorted and converted to .bam files:

> (samtools view -S -b out.sam > out.bam; rm -rf out.sam out.sorted.sam; samtools sort out.bam -o out.sorted.bam; rm -rf out.bam).

TIMEOR plots the percent of reads mapping exactly once and overall alignment rate, which highlights that replicate SRR8843750_1.fastq.gz (timepoint 40 minutes) and replicate SRR8843747_1.fastq.gz (timepoint 180 minutes) show low alignment (**Figure S4A, B**). Due to contamination at the experimental level, replicate SRR8843750_1.fastq.gz was removed from further analysis. Next all replicates were converted to sorted .bam files (Li et al., 2009) to produce read counts per gene using HTSeq (Anders et al., 2015). Note that htseq assumes that .bam is sorted by name by default and not position, so -r pos is necessary if samtools sort is used without the -n flag. TIMEOR ran HTSeq to produce a read count matrix of samples by read counts per gene for each sample, and TIMEOR then merged the samples in chronological order to create the final read count matrix of genes by sample (i.e. replicates). Specifically, the used command for HTSeq is:

> (htseq-count -f bam -r pos -i gene_id out.sorted.bam BDGP6/genes.gtf > htseq_counts).

##### Removing contamination

Note that replicate SRR8843750 shows an overall mapping rate of 17.44%. Further, SRR8843747 shows an overall mapping rate of 47.02%. When we use BLAST (both blastn and megablast, Altschul et al., 1990) to identify several sequences from replicate SRR8843750 we see synthetic construct external RNA control and Methanocaldococcus jannaschii as the top subject sequence. When we use BLAST to identify several sequences from replicate SRR8843747 we see *Drosophila simulans* uncharacterized protein as the top subject sequence. In contrast, looking at several sequences from replicate SRR8843757 and SRR8843749 the top subject sequences are derived from *Drosophila melanogaster*. Therefore, the SRR8843750 replicate was removed.

##### Normalization and correction

After filtering out rows where the mean read count is 5 or less across all replicates and samples, 8909/17558 genes remained. This normalization and correction was used for Next maSigPro (Nueda et al., 2014) as it requires pre-normalization and correction. See further explanation in section “Next maSigPro results”.

##### Differential expression

According to comparisons made by Spies et al., 2019 (Spies et al., 2019): all time course tools were outperformed by classical pairwise comparison approaches (in this case DESeq2, Love et al., 2014) on short time series (<8 time points) in terms of overall performance and robustness to noise, mostly because of high number of false positives, with the exception of ImpulseDE2 (Fischer et al., 2018). On longer time series, pairwise approaches were performed less well than splineTimeR (Michna et al., 2016) and Next maSigPro, which did not identify any false-positive candidates. SplineTimeR does not support Zirin et al. 2019’s experimental design and thus was unable to be run. Other methods consider this dataset experimental set-up within their modeling framework.

There is significant overlap between the differentially expressed (DE) genes previously called by Zirin et al., 2019 and these three methods. ImpulseDE2 (Fischer et al., 2018) showed the most significant set of DE genes overlapping with the previous study (Zirin et al., 2019). This gene set was used downstream in TIMEOR to produce a clustermap of seven gene trajectory clusters. We recapitulate the findings of Zirin et al., 2019 by finding two clusters of DE genes (adjusted p-value < 0.05) enriched in the GO categories nucleolus and ribosome biogenesis, some of which were not identified previously. TIMEOR identified several additional clusters with novel characteristics.

Zirin et al., 2019 identified 1211 DE genes and followed-up to identify a subset of 33 DE genes involved in several biological processes including ribosome biogenesis and transcription. Converting those gene symbols to Flybase IDs gives 36 genes, as some gene names correspond to one of several Flybase IDs. Comparing the gene list of 36 Flybase IDs between all methods, we observe the most significant gene overlap with ImpulseDE2 (p-value < 5.518766e-08 using the hypergeometric test of significance) compared to Next maSigPro (p-value <0.0008382603) and DESeq2 (p-value <0.002140549). Comparing our results with the 1211 DE genes from Zirin et al.,TIMEOR finds that ImpulseDE2 showed the highest overlap with the list of 1211 DE genes (p-value = 5.33342e-127 using the hypergeometric test of significance) and the highest overlap with other methods (**Figure S4C**).

##### ImpulseDE2 results

ImpulseDE2 requires unnormalized and uncorrected counts and replicate data. At an adjusted p-value cutoff of 0.05, we found 156 significantly differentially expressed genes. As aforementioned in the previous paragraph, ImpulseDE2 shows the most significant gene overlap with the previous study when looking at the full set of 1211 DE genes (p-value = 5.33342e-127) and the followup subset of 36 DE genes (p-value < 5.518766e-08).

##### Next maSigPro results

Data must be normalized before the application of Next maSigPro because there is no integrated normalization method (Nueda et al., 2014). Thus, the data are first filtered to remove any genes across all replicates with less than a mean of 5 reads. The data are then normalized by the mean ratio normalization and batch effect corrected with Combat (Leek et al., 2012). Considering all case conditions with 3 batches, DESeq2 reported 76 differentially expressed genes. When comparing with the full set of 1211 DE genes there is a less significant overlap (p-value = 3.748126e-69) than with ImpulseDE2. The overlap between these 76 genes and the 156 from ImpulseDE2 is 76 genes. Furthermore, when comparing with the follow-up subset of 36 DE genes, Next maSigPro reports a less significant overlap with Zirin et al., 2019 than ImpulseDE2, with 3 overlapping genes, resulting in a p-value of 0.0006961443.

##### DESeq2 results

DESeq2 requires unnormalized and uncorrected counts with replicate data. Given that this is a categorical differential expression method, we must test for any differences over multiple time points all compared to time point 0 (i.e. the reference). Here we use a design including the factor of time, and use the likelihood ratio test to see if there are significant changes in expression for genes between T0 and the each additional time point. Considering all case conditions with 3 batches, we obtain 30 differentially expressed genes. When comparing with the full set of 1211 DE genes there is the least significant overlap (p-value = 2.010729e-29) with Zirin et al., 2019. Furthermore, when comparing with the follow-up subset of 36 DE genes, DESeq2 reports the least significant overlap with 3 overlapping genes, resulting in a p-value of 4.322579e-05.

##### TIMEOR clustermap of z-scored gene expression trajectories at an adjusted p-value<0.05

**Figure 2B and 3A** show the clustermap of DEG expression as z-scores, which correspond to the observed read counts of a sample rescaled to a standard normal distribution (with mean 0 and standard deviation 1). Specifically, the z-score of an observation is the mean divided by the standard deviation of all observations in the sample. Thus, the z-score describes how many standard deviations an observation is from the mean of all observations, with positive values indicating upregulation of the gene and negative values indicating downregulation of a gene. These z-scored DEG trajectories were then clustered using Euclidean distance and Ward.D2 (Murtagh et al., 2014) hierarchical clustering. Note that z-scored expression values are generated for ImpulseDE2 and Next maSigPro. DESeq2 is a categorical method that outputs log2 fold change values across each timepoint. TIMEOR uses a variety of methods to assess stability of the number of clusters. In particular, the Elbow method showed that the decreasing error stabilized at six clusters (**Figure S4D**). Further human inspection showed a clear delineation of six clusters in the heatmap.

##### TIMEOR GO analysis

ClusterProfiler (Yu et al., 2012) is run to determine gene ontology (GO) enrichment of each cluster of genes (**Figure 2B and 3A**) within the categories of biological process, cellular component, and molecular function. Within each of our six clusters using a Benjamini Hochberg adjusted p-value cutoff of 0.05, TIMEOR found the most significant GO term for cluster d to be cellular response to endogenous stimulus within “biological process”. In cluster c, the nucleolus (cellular component), ribosome biogenesis (biological process) and catalytic activity acting on RNA (molecular function) are the most enriched terms. Within the largest cluster b, ribosome biogenesis (biological process), nucleolus (cellular component) and snoRNA binding (molecular function) are the most enriched terms. The molecular function enrichment for snoRNA binding is particularly interesting because in the next activated cluster e there is a significant number of snoRNAs present (**Figure 2E**). Our full TIMEOR results folder with various GO analysis visualizations can be found on Github.

##### TIMEOR pathway analysis

Pathview (Luo et al. 2013) is run to highlight genes and their trajectories (−1 is most downregulated and 1 is most upregulated) in any enriched pathways for each cluster of genes. **Figure 2D** shows the DE genes in cluster b which are part of the ribosome biogenesis pathway. This pathway was also identified in the previous study which generated these data from Zirin et al., 2019.

##### TIMEOR factor binding analysis

For each cluster, TIMEOR found both observed and predicted TFs via the *Top Predicted Transcription Factors by Orthology* table and *Top Predicted Transcription Factors by Motif Similarity* table, and each individual method’s results which are visible in their associated *Top Transcription Factors per Method Table* for both orthology motif similarity. In TIMEOR’s factor binding script, the threshold normalized enrichment score was set to 3, and the percent concordance threshold was set to 30. **Figure 3B** shows the prioritized list of TFs per cluster. TIMEOR’s provided list of ENCODE IDs (Davis et al., 2018; Joly Beauparlant et al., 2020) enabled us to choose which factor data to upload and view via average profiles. See next section for those details.

##### TIMEOR average profile generation

For each top predicted transcription factor, we identified the most closely related ChIP-seq dataset (in .bigwig format) on ENCODE using the IDs provided by TIMEOR (**Supplementary Data 2**). Specifically, using data from Dr. Kevin White, we used 6-24 whole organism embryo ChIP-seq data for CWO (ENCFF491LHJ). We used 0-16 hours whole organism embryo ChIP-seq data for MYC (ENCFF829HXS). We used whole organism prepupa ChIP-seq data for CYC (ENCFF082KKV). And we used whole organism wandering third instar larva for HNF4 (ENCFF680FFM), TBP (ENCFF553PBY), and CG9727 (ENCFF145ATU). TIMEOR then uses deepTools (Ramírez et al., 2016) to plot the average peak distribution over each gene trajectory cluster, specifically using the set of commands below:

> (computeMatrix scale-regions -S ${INPUT_BIGWIG} -R ${LIST_BEDS[@]} --binSize 250--beforeRegionStartLength 1000 --afterRegionStartLength 1000 --regionBodyLength 5000 -o ${OUTPUT_DIR}/matrix.genes.clusters.mat.gz --skipZeros --smartLabels --sortRegions descend)

> (plotHeatmap -m ${OUTPUT_DIR}/matrix.genes.clusters.mat.gz -out ${OUTPUT_DIR}/heatmap_genes.clusters.${TF}.png --heatmapHeight 25 --heatmapWidth 15--labelRotation 45 --missingDataColor red --whatToShow=“heatmap only”)

> (plotProfile -m ${OUTPUT_DIR}/matrix.genes.clusters.mat.gz -out ${OUTPUT_DIR}/avg_profile.${TF}.png --plotType=se --labelRotation 45 --plotHeight 15 --plotWidth 15 --colors ${COLORS})

##### TIMEOR temporal relations

Between all clusters, TIMEOR highlighted the temporal relationships between all observed and predicted TFs (those listed by default in the *Top Predicted Transcription Factors by Orthology* table). In TIMEOR’s temporal relations script, the number of time steps over which one gene can influence the transcription of another gene (question six) was set to two. Within this time window, all possible directed interactions are considered and characterized. We used the last tab of Secondary Analysis (i.e. “Temporal Relations”) to visualize and interpret the temporal relations table (encoding the GRN) in combination with the transcription factor network (Franceschini et al., 2013), the clustermap, and the list of predicted TFs to identify the main altered regulatory mechanisms. Overall, TIMEOR predicts a GRN in which first, predicted TF Hnf4 is predicted to repress observed gene *CG32772. CG32772* is then predicted to activate the observed gene *cwo*. Predicted TF TBP is also predicted to activate *cwo*. Due to TBP’s poor binding profile with its predicted cluster and overall, we examined CYC (**Figure 3B, S4F**) and determined that it is likely activating *cwo* (**Figure 3D**). Observed TF CWO is then predicted to activate *pdp1*.

### RESOURCE AVAILABILITY

The code generated during this study is available on Github at https://github.com/ashleymaeconard/TIMEOR.git.

- The published article includes all code generated or analyzed during this study.
- This study reanalysed data generated and available on GEO with accession numbers GSE118691, and SRP190499. This study also reanalyzed the following ChIP-seq data ENCFF491LHJ, ENCFF829HXS, ENCFF082KKV, ENCFF680FFM, ENCFF553PBY, and ENCFF145ATU.

## Notes

https://github.com/ashleymaeconard/TIMEOR

### Competing Interest Statement

The authors have declared no competing interest.

## Bibliography

Afgan, E., Baker, D., Batut, B., van den Beek, M., Bouvier, D., Cech, M., Chilton, J., Clements, D., Coraor, N., Grüning, B.A., et al. (2018). The Galaxy platform for accessible, reproducible and collaborative biomedical analyses: 2018 update. Nucleic Acids Res. 46, W537–W544.

Aibar, S., González-Blas, C.B., Moerman, T., Huynh-Thu, V.A., Imrichova, H., Hulselmans, G., Rambow, F., Marine, J.-C., Geurts, P., Aerts, J., et al. (2017). SCENIC: single-cell regulatory network inference and clustering. Nat. Methods 14, 1083–1086.

Altschul, S. F., Gish, W., Miller, W., Myers, E. W., & Lipman, D. J. (1990). Basic local alignment search tool. Journal of molecular biology, 215(3), 403–410.

Anders, S., Pyl, P. T., & Huber, W. (2015). HTSeq—a Python framework to work with high-throughput sequencing data. Bioinformatics, 31(2), 166–169.

Andrews, S. (2010). FastQC: a quality control tool for high throughput sequence data.

Bailey, T.L., and Elkan, C. (1994). Fitting a mixture model by expectation maximization to discover motifs in biopolymers. Proc. Int. Conf. Intell. Syst. Mol. Biol. 2, 28–36.

Bansal, M., Della Gatta, G., and di Bernardo, D. (2006). Inference of gene regulatory networks and compound mode of action from time course gene expression profiles. Bioinformatics 22, 815–822.

Bar-Joseph, Z., Gerber, G.K., Lee, T.I., Rinaldi, N.J., Yoo, J.Y., Robert, F., Gordon, D.B., Fraenkel, E., Jaakkola, T.S., Young, R.A., et al. (2003). Computational discovery of gene modules and regulatory networks. Nat. Biotechnol. 21, 1337–1342.

Barbosa, S., Niebel, B., Wolf, S., Mauch, K., and Takors, R. (2018). A guide to gene regulatory network inference for obtaining predictive solutions: Underlying assumptions and fundamental biological and data constraints. BioSystems 174, 37–48.

Barry, W.E., and Thummel, C.S. (2016). The Drosophila HNF4 nuclear receptor promotes glucose-stimulated insulin secretion and mitochondrial function in adults. Elife 5.

Berk, A.J. (2016). Discovery of RNA splicing and genes in pieces. Proc Natl Acad Sci USA 113, 801–805.

Boden, G., Ruiz, J., Urbain, J.L., and Chen, X. (1996). Evidence for a circadian rhythm of insulin secretion. Am. J. Physiol. 271, E246–52.

Bolger, A. M., Lohse, M., & Usadel, B. (2014). Trimmomatic: A flexible trimmer for Illumina Sequence Data. Bioinformatics, btu170.

Brent, M.R. (2016). Past roadblocks and new opportunities in transcription factor network mapping. Trends Genet. 32, 736–750.

Caliński, T., & Harabasz, J. (1974). A dendrite method for cluster analysis. Communications in Statistics-theory and Methods, 3(1), 1–27.

Chai, L. E., Loh, S. K., Low, S. T., Mohamad, M. S., Deris, S., & Zakaria, Z. (2014). A review on the computational approaches for gene regulatory network construction. Computers in biology and medicine, 48, 55–65.

Chang W., Cheng J., Allaire J.J., Xie Y., and McPherson, J. (2020). shiny: Web Application Framework for R. R package version 1.4.0.2. https://CRAN.R-project.org/package=shiny

Cornwell, M., Vangala, M., Taing, L., Herbert, Z., Köster, J., Li, B., Sun, H., Li, T., Zhang, J., Qiu, X., et al. (2018). VIPER: Visualization Pipeline for RNA-seq, a Snakemake workflow for efficient and complete RNA-seq analysis. BMC Bioinformatics 19, 135.

Davis, C.A., Hitz, B.C., Sloan, C.A., Chan, E.T., Davidson, J.M., Gabdank, I., Hilton, J.A., Jain, K., Baymuradov, U.K., Narayanan, A.K., et al. (2018). The Encyclopedia of DNA elements (ENCODE): data portal update. Nucleic Acids Res. 46, D794–D801.

Edgar, R., Domrachev, M., and Lash, A.E. (2002). Gene Expression Omnibus: NCBI gene expression and hybridization array data repository. Nucleic Acids Res. 30, 207–210.

Ewels, P., Magnusson, M., Lundin, S., & Käller, M. (2016). MultiQC: summarize analysis results for multiple tools and samples in a single report. Bioinformatics, 32(19), 3047–3048.

Faith, J.J., Hayete, B., Thaden, J.T., Mogno, I., Wierzbowski, J., Cottarel, G., Kasif, S., Collins, J.J., and Gardner, T.S. (2007). Large-scale mapping and validation of Escherichia coli transcriptional regulation from a compendium of expression profiles. PLoS Biol. 5, e8.

Fathallah-Shaykh, H.M. (2010). Dynamics of the Drosophila circadian clock: theoretical anti-jitter network and controlled chaos. PLoS ONE 5, e11207.

Fischer, D.S., Theis, F.J., and Yosef, N. (2018). Impulse model-based differential expression analysis of time course sequencing data. Nucleic Acids Res. 46, e119.

Franceschini, A., Szklarczyk, D., Frankild, S., Kuhn, M., Simonovic, M., Roth, A., Lin, J., Minguez, P., Bork, P., von Mering, C., et al. (2013). STRING v9.1: protein-protein interaction networks, with increased coverage and integration. Nucleic Acids Res. 41, D808–15.

Frazee, A. C., Jaffe, A. E., Langmead, B., & Leek, J. T. (2015). Polyester: simulating RNA-seq datasets with differential transcript expression. Bioinformatics, 31(17), 2778–2784.

Gale, J.E., Cox, H.I., Qian, J., Block, G.D., Colwell, C.S., and Matveyenko, A.V. (2011). Disruption of circadian rhythms accelerates development of diabetes through pancreatic beta-cell loss and dysfunction. J. Biol. Rhythms 26, 423–433.

Ge, S.X., Son, E.W., and Yao, R. (2018). iDEP: an integrated web application for differential expression and pathway analysis of RNA-Seq data. BMC Bioinformatics 19, 534.

Granger, C. W. (1969). Investigating causal relations by econometric models and cross-spectral methods. Econometrica: journal of the Econometric Society, 424–438.

Han, H., Cho, J.-W., Lee, S., Yun, A., Kim, H., Bae, D., Yang, S., Kim, C.Y., Lee, M., Kim, E., et al. (2018). TRRUST v2: an expanded reference database of human and mouse transcriptional regulatory interactions. Nucleic Acids Res. 46, D380–D386.

Haury, A.-C., Mordelet, F., Vera-Licona, P., and Vert, J.-P. (2012). TIGRESS: Trustful Inference of Gene REgulation using Stability Selection. BMC Syst. Biol. 6, 145.

Heinz, S., Benner, C., Spann, N., Bertolino, E., Lin, Y. C., Laslo, P.… & Glass, C. K. (2010). Simple combinations of lineage-determining transcription factors prime cis-regulatory elements required for macrophage and B cell identities. Molecular cell, 38(4), 576–589.

Huynh-Thu, V.A., and Geurts, P. (2018). dynGENIE3: dynamical GENIE3 for the inference of gene networks from time series expression data. Sci. Rep. 8, 3384.

Huynh-Thu, V.A., Irrthum, A., Wehenkel, L., and Geurts, P. (2010). Inferring regulatory networks from expression data using tree-based methods. PLoS ONE 5.

James, S.M., Honn, K.A., Gaddameedhi, S., and Van Dongen, H.P.A. (2017). Shift Work: Disrupted Circadian Rhythms and Sleep-Implications for Health and Well-Being. Curr. Sleep Med. Rep. 3, 104–112.

Jensen, T.L., Frasketi, M., Conway, K., Villarroel, L., Hill, H., Krampis, K., and Goll, J.B. (2017). RSEQREP: RNA-Seq Reports, an open-source cloud-enabled framework for reproducible RNA-Seq data processing, analysis, and result reporting. [version 2; peer review: 2 approved]. F1000Res. 6, 2162.

Johnson, D.S., Mortazavi, A., Myers, R.M., and Wold, B. (2007). Genome-wide mapping of in vivo protein-DNA interactions. Science 316, 1497–1502.

Joly Beauparlant C., Lemacon A., Fournier E., Droit A. (2020). ENCODExplorer: A compilation of ENCODE metadata. R package version 2.14.0.

de Jong, A., van der Meulen, S., Kuipers, O.P., and Kok, J. (2015). T-REx: Transcriptome analysis webserver for RNA-seq Expression data. BMC Genomics 16, 663.

Kadener, S., Stoleru, D., McDonald, M., Nawathean, P., and Rosbash, M. (2007). Clockwork Orange is a transcriptional repressor and a new Drosophila circadian pacemaker component. Genes Dev. 21, 1675–1686.

Kang, W., Jia, Z., Tang, D., Zhao, X., Shi, J., Jia, Q., He, K., and Feng, Q. (2019a). Time-Course Transcriptome Analysis for Drug Repositioning in Fusobacterium nucleatum-Infected Human Gingival Fibroblasts. Front. Cell Dev. Biol. 7, 204.

Kang, W., Jia, Z., Tang, D., Zhang, Z., Gao, H., He, K., and Feng, Q. (2019b). Fusobacterium nucleatum Facilitates Apoptosis, ROS Generation, and Inflammatory Cytokine Production by Activating AKT/MAPK and NF-κB Signaling Pathways in Human Gingival Fibroblasts. Oxid. Med. Cell. Longev. 2019, 1681972.

Kang, Y., Liow, H.-H., Maier, E.J., and Brent, M.R. (2018). NetProphet 2.0: mapping transcription factor networks by exploiting scalable data resources. Bioinformatics 34, 249–257.

Kartashov, A.V., and Barski, A. (2015). BioWardrobe: an integrated platform for analysis of epigenomics and transcriptomics data. Genome Biol. 16, 158.

Kaufman, L., & Rousseeuw, P. J. (2009). Finding groups in data: an introduction to cluster analysis (Vol. 344). John Wiley & Sons.

Khan, A., & Mathelier, A. (2017). Intervene: a tool for intersection and visualization of multiple gene or genomic region sets. BMC bioinformatics, 18(1), 287.

Kim, D., Paggi, J. M., Park, C., Bennett, C., & Salzberg, S. L. (2019). Graph-based genome alignment and genotyping with HISAT2 and HISAT-genotype. Nature biotechnology, 37(8), 907–915.

Křivánek, M., & Morávek, J. (1986). NP-hard problems in hierarchical-tree clustering. Acta informatica, 23(3), 311–323.

Kulakovskiy, I. V., Vorontsov, I. E., Yevshin, I. S., Sharipov, R. N., Fedorova, A. D., Rumynskiy, E. I.… & Kolpakov, F. A. (2018). HOCOMOCO: towards a complete collection of transcription factor binding models for human and mouse via large-scale ChIP-Seq analysis. Nucleic acids research, 46(D1), D252–D259.

Langfelder, P., and Horvath, S. (2008). WGCNA: an R package for weighted correlation network analysis. BMC Bioinformatics 9, 559.

Langmead, B., & Salzberg, S. L. (2012). Fast gapped-read alignment with Bowtie 2. Nature methods, 9(4), 357.

Leek, J. T., Johnson, W. E., Parker, H. S., Jaffe, A. E., & Storey, J. D. (2012). The sva package for removing batch effects and other unwanted variation in high-throughput experiments. Bioinformatics, 28(6), 882–883.

Leinonen, R., Sugawara, H., Shumway, M., & International Nucleotide Sequence Database Collaboration. (2010). The sequence read archive. Nucleic acids research, 39(suppl_1), D19–D21.

Liu, W., Zhu, W., Liao, B., and Chen, X. (2016). Gene Regulatory Network Inferences Using a Maximum Relevance and Maximum-Significance Strategy. PLoS ONE 11, e0166115.

Love, M.I., Huber, W., and Anders, S. (2014). Moderated estimation of fold change and dispersion for RNA-seq data with DESeq2. Genome Biol. 15, 550.

Luo, W., and Brouwer, C. (2013). Pathview: an R/Bioconductor package for pathway-based data integration and visualization. Bioinformatics 29, 1830–1831.

Marbach, D., Costello, J.C., Küffner, R., Vega, N.M., Prill, R.J., Camacho, D.M., Allison, K.R., DREAM5 Consortium, Kellis, M., Collins, J.J., et al. (2012). Wisdom of crowds for robust gene network inference. Nat. Methods 9, 796–804.

Margolin, A.A., Nemenman, I., Basso, K., Wiggins, C., Stolovitzky, G., Dalla Favera, R., and Califano, A. (2006). ARACNE: an algorithm for the reconstruction of gene regulatory networks in a mammalian cellular context. BMC Bioinformatics 7 Suppl 1, S7.

Maury, E. (2019). Off the clock: from circadian disruption to metabolic disease. Int. J. Mol. Sci. 20.

Michna, A., Braselmann, H., Selmansberger, M., Dietz, A., Hess, J., Gomolka, M., Hornhardt, S., Blüthgen, N., Zitzelsberger, H., and Unger, K. (2016). Natural Cubic Spline Regression Modeling Followed by Dynamic Network Reconstruction for the Identification of Radiation-Sensitivity Gene Association Networks from Time-Course Transcriptome Data. PLoS ONE 11, e0160791.

Mochida, K., Koda, S., Inoue, K., and Nishii, R. (2018). Statistical and machine learning approaches to predict gene regulatory networks from transcriptome data sets. Front. Plant Sci. 9, 1770.

Murtagh, F., & Legendre, P. (2014). Ward’s hierarchical agglomerative clustering method: which algorithms implement Ward’s criterion?. Journal of classification, 31(3), 274–295.

Nueda, M.J., Tarazona, S., and Conesa, A. (2014). Next maSigPro: updating maSigPro bioconductor package for RNA-seq time series. Bioinformatics 30, 2598–2602.

Oytam, Y., Sobhanmanesh, F., Duesing, K., Bowden, J. C., Osmond-McLeod, M., & Ross, J. (2016). Risk-conscious correction of batch effects: maximising information extraction from high-throughput genomic datasets. BMC bioinformatics, 17(1), 332.

Qian, J., and Scheer, F.A.J.L. (2016). Circadian system and glucose metabolism: implications for physiology and disease. Trends Endocrinol. Metab. 27, 282–293.

Radovic, M., Ghalwash, M., Filipovic, N., and Obradovic, Z. (2017). Minimum redundancy maximum relevance feature selection approach for temporal gene expression data. BMC Bioinformatics 18, 9.

Ramírez, F., Ryan, D. P., Grüning, B., Bhardwaj, V., Kilpert, F., Richter, A. S.… & Manke, T. (2016). deepTools2: a next generation web server for deep-sequencing data analysis. Nucleic acids research, 44(W1), W160–W165.

Reynolds, A. P., Richards, G., de la Iglesia, B., & Rayward-Smith, V. J. (2006). Clustering rules: a comparison of partitioning and hierarchical clustering algorithms. Journal of Mathematical Modelling and Algorithms, 5(4), 475–504.

Rousseeuw, P. J. (1987). Silhouettes: a graphical aid to the interpretation and validation of cluster analysis. Journal of computational and applied mathematics, 20, 53–65.

Roy, S., Lagree, S., Hou, Z., Thomson, J.A., Stewart, R., and Gasch, A.P. (2013). Integrated module and gene-specific regulatory inference implicates upstream signaling networks. PLoS Comput. Biol. 9, e1003252.

Sahraeian, S.M.E., Mohiyuddin, M., Sebra, R., Tilgner, H., Afshar, P.T., Au, K.F., Bani Asadi, N., Gerstein, M.B., Wong, W.H., Snyder, M.P., et al. (2017). Gaining comprehensive biological insight into the transcriptome by performing a broad-spectrum RNA-seq analysis. Nat. Commun. 8, 59.

Sandelin, A., Alkema, W., Engström, P., Wasserman, W. W., & Lenhard, B. (2004). JASPAR: an open access database for eukaryotic transcription factor binding profiles. Nucleic acids research, 32(suppl_1), D91–D94.

Sharp, P.A. (1991). “Five easy pieces”. Science 254, 663.

Sharp, P.A. (2009). The centrality of RNA. Cell 136, 577–580.

Sharp, B., Paquet, E., Naef, F., Bafna, A., and Wijnen, H. (2017). A new promoter element associated with daily time keeping in Drosophila. Nucleic Acids Res. 45, 6459–6470.

Sievert C (2020). Interactive Web-Based Data Visualization with R, plotly, and shiny. Chapman and Hall/CRC. ISBN 9781138331457, https://plotly-r.com.

Skene, P.J., and Henikoff, S. (2017). An efficient targeted nuclease strategy for high-resolution mapping of DNA binding sites. Elife 6.

Spies, D., and Ciaudo, C. (2015). Dynamics in Transcriptomics: Advancements in RNA-seq Time Course and Downstream Analysis. Comput. Struct. Biotechnol. J. 13, 469–477.

Spies, D., Renz, P.F., Beyer, T.A., and Ciaudo, C. (2019). Comparative analysis of differential gene expression tools for RNA sequencing time course data. Brief. Bioinformatics 20, 288–298.

Spurney, R.J., Van den Broeck, L., Clark, N.M., Fisher, A.P., de Luis Balaguer, M.A., and Sozzani, R. (2020). tuxnet: a simple interface to process RNA sequencing data and infer gene regulatory networks. Plant J. 101, 716–730.

Stenvers, D.J., Scheer, F.A.J.L., Schrauwen, P., la Fleur, S.E., and Kalsbeek, A. (2019). Circadian clocks and insulin resistance. Nat. Rev. Endocrinol. 15, 75–89.

Szklarczyk, D., Gable, A. L., Lyon, D., Junge, A., Wyder, S., Huerta-Cepas, J., Simonovic, M., Doncheva, N. T., Morris, J. H., Bork, P., Jensen, L. J., & Mering, C. V. (2019). STRING v11: protein-protein association networks with increased coverage, supporting functional discovery in genome-wide experimental datasets. Nucleic acids research, 47(D1), D607–D613. https://doi.org/10.1093/nar/gky1131

Torre, D., Lachmann, A., and Ma’ayan, A. (2018). BioJupies: Automated Generation of Interactive Notebooks for RNA-Seq Data Analysis in the Cloud. Cell Syst. 7, 556–561.e3.

Thieurmel, B. (2016) VisNetwork. https://www.rdocumentation.org/packages/visNetwork/versions/2.0.1. Accessed 12 Dec 2019.

Verfaillie, A., Imrichová, H., Van de Sande, B., Standaert, L., Christiaens, V., Hulselmans, G.… & Atak, Z. K. (2014). iRegulon: from a gene list to a gene regulatory network using large motif and track collections. PLoS Comput Biol, 10(7), e1003731.

Villaverde, A.F., Ross, J., Morán, F., and Banga, J.R. (2014). MIDER: network inference with mutual information distance and entropy reduction. PLoS ONE 9, e96732.

Wang, Y.-H., and Huang, M.-L. (2009). Reduction of Lobe leads to TORC1 hypoactivation that induces ectopic Jak/STAT signaling to impair Drosophila eye development. Mech. Dev. 126, 781–790.

Wang, Y.K., Hurley, D.G., Schnell, S., Print, C.G., and Crampin, E.J. (2013). Integration of steady-state and temporal gene expression data for the inference of gene regulatory networks. PLoS ONE 8, e72103.

Wang, Z., Gerstein, M., and Snyder, M. (2009). RNA-Seq: a revolutionary tool for transcriptomics. Nat. Rev. Genet. 10, 57–63.

Wani, N., and Raza, K. (2019). Integrative approaches to reconstruct regulatory networks from multi-omics data: A review of state-of-the-art methods. Comput. Biol. Chem. 83, 107120.

Weirauch, M. T., Yang, A., Albu, M., Cote, A. G., Montenegro-Montero, A., Drewe, P.… & Zheng, H. (2014). Determination and inference of eukaryotic transcription factor sequence specificity. Cell, 158(6), 1431–1443.

Wilkinson, A.C., Nakauchi, H., and Göttgens, B. (2017). Mammalian transcription factor networks: recent advances in interrogating biological complexity. Cell Syst. 5, 319–331.

Wingender, E., Dietze, P., Karas, H., & Knüppel, R. (1996). TRANSFAC: a database on transcription factors and their DNA binding sites. Nucleic acids research, 24(1), 238–241.

Yu, G., Wang, L. G., Han, Y., & He, Q. Y. (2012). clusterProfiler: an R package for comparing biological themes among gene clusters. Omics: a journal of integrative biology, 16(5), 284–287.

Zhang, L., Feng, X.K., Ng, Y.K., and Li, S.C. (2016). Reconstructing directed gene regulatory network by only gene expression data. BMC Genomics 17 Suppl 4, 430.

Zhou, J., Yu, W., and Hardin, P.E. (2016). CLOCKWORK ORANGE Enhances PERIOD Mediated Rhythms in Transcriptional Repression by Antagonizing E-box Binding by CLOCK-CYCLE. PLoS Genet. 12, e1006430.

Zhu, L. J., Christensen, R. G., Kazemian, M., Hull, C. J., Enuameh, M. S., Basciotta, M. D.… & Sinha, S. (2011). FlyFactorSurvey: a database of Drosophila transcription factor binding specificities determined using the bacterial one-hybrid system. Nucleic acids research, 39(suppl_1), D111–D117.

Zirin, J., Ni, X., Sack, L.M., Yang-Zhou, D., Hu, Y., Brathwaite, R., Bulyk, M.L., Elledge, S.J., and Perrimon, N. (2019). Interspecies analysis of MYC targets identifies tRNA synthetases as mediators of growth and survival in MYC-overexpressing cells. Proc Natl Acad Sci USA 116, 14614–14619.

Zoppoli, P., Morganella, S., and Ceccarelli, M. (2010). TimeDelay-ARACNE: Reverse engineering of gene networks from time-course data by an information theoretic approach. BMC Bioinformatics 11, 154.

